# Multimodal brain cell atlas across the adult macaque lifespan

**DOI:** 10.1101/2025.03.11.641786

**Authors:** Xiao Zhang, Guangyao Lai, Xiangyu Guo, Wen Ma, Yue Yuan, Han Zhang, Peng Fan, Xingyuan Liu, Pengcheng Guo, Juan An, Jinxiu Li, Keke Huang, Jing Zuo, Liang Wu, Shuncheng Shangguan, Feng Yu, Yuchen Xiang, Yuebei Hu, Yunting Huang, Yiting Lin, Dajian He, Mingtian Pan, Laiqiang Chen, Bang Li, Weili Yang, Peng Yin, Shihua Li, Xiaolan Zhang, Tao Yang, Jing Chen, Qiaoling Zhong, Yuan Lv, Baoming Qin, Xiaobing Qing, Qiuting Deng, Yaling Huang, Chang Liu, Lei Han, Shiping Liu, Dajiang Qin, Zhen Liu, Yi Zhou, Liang Qiao, Chengyu Li, YanXiao Zhang, Giacomo Volpe, Simon Graf, Ian Ferenc Diks, Matthias Flotho, Rafal P. Krol, Jan Mulder, Andrew P. Hutchins, Andreas Keller, Patrick H. Maxwell, David C. Rubinsztein, Xiaolei Liu, Hung-Fat Tse, Muming Poo, Bo Wang, Longqi Liu, Ying Gu, Jun Xie, Xun Xu, Xiaojiang Li, Chuanyu Liu, Yiwei Lai, Mingyuan Liu, Miguel A. Esteban

## Abstract

High-throughput single-cell omics of non-human primate tissues present a remarkable opportunity to study primate brain aging. Here, we introduce a transcriptomic and chromatin accessibility landscape of 1,985,317 cells from eight brain regions of 13 cynomolgus female monkeys spanning adult lifespan including exceptionally old individuals up to 29-years old. This dataset uncovers dynamic molecular changes in critical brain functions such as synaptic communication and axon myelination, exhibiting a high degree of cell type and brain region specificity. We identify the multicellular networks of the pons and medulla as a previously unrecognized hotspot for aging. Furthermore, comparative analyses with human neurodegeneration datasets highlight both shared and distinct mechanisms contributing to aging and disease. In addition, we uncover transcription factors implicated in monkey brain aging and pinpoint aging-regulated loci linked to longevity and neurodegeneration. This spatiotemporal atlas will advance our understanding of primate brain aging and its broader implications for health and disease.

## INTRODUCTION

The mammalian brain is composed of diverse neuronal and non-neuronal cell types distributed across regions derived ontogenetically from distinct neuroectodermal and non-neuroectodermal domains^1–3^. This cellular diversity arises through the activation of specific genetic programs influenced by local signaling cues, long-range intercellular interactions, and cell migrations. The resulting regional microenvironments not only shape brain functions but also impose region– and cell-type-specific responses to stress and injury. As life progresses, regional multicellular networks undergo progressive alterations, leading to impairments in critical brain functions such as synaptic communication, axon myelination, and cell viability. These changes contribute to deficits in learning and memory, attention, decision-making, sensory perception, and motor abilities. The disruption of multicellular networks is further exacerbated in aging-related neurodegenerative diseases such as Alzheimer’s disease (AD)^4^, the most prevalent neurodegenerative disorder in older adults and a significant global socioeconomic health challenge. Extensive research has been conducted, yet the extent to which brain aging and neurodegeneration share molecular mechanisms and impact the same cell types and regions remains poorly understood. Addressing these questions is crucial for improving brain health in aging populations and may uncover therapeutic opportunities for AD. Moreover, given the brain’s integral role in regulating most bodily functions, enhancing brain health is also essential for promoting overall longevity and quality of life.

Over the past few decades, significant efforts have been dedicated to understanding normal and pathological brain functions across various mammalian species, with a primary focus on rodents^5–8^. However, primate brains exhibit distinct anatomical and molecular characteristics compared to those of other mammals^9^. The advent of high-throughput single-cell omics has recently advanced the study of primate brain aging and neurodegeneration^4,10–13^. Despite these achievements, systematic studies in humans face several challenges, including high interindividual heterogeneity, the difficulty of obtaining timely postmortem samples, and limitations in sample storage and availability in brain banks. Non-human primates (NHPs) offer a valuable alternative for studying human brain aging, owing to their evolutionary proximity and physiological similarities to our species. Additionally, farmed NHPs exhibit reduced biological and lifestyle variability, providing a more controlled model for investigating brain aging and its associated processes^14^.

While recent studies have investigated the effects of aging in selected regions of the NHP brain^10,11,15,16^ (**Table S1**), no study to date has comprehensively analyzed multiple regions across individuals spanning a full range of ages, from young adulthood to exceptionally old age. Analyzing multiple regions from the same individuals is essential for identifying both general and region-specific vulnerabilities in a comparative manner. Additionally, including exceptionally old individuals may help uncover the molecular drivers of healthy brain aging and, potentially, longevity. Furthermore, epigenetic data on aging in the NHP brain, particularly insights gained from accessible chromatin analysis, remain largely unavailable. Such information could provide valuable understanding of the gene regulatory mechanisms underlying primate brain aging.

Here, we have constructed a multimodal NHP aging brain cell atlas (NHPABC) combining single-nucleus RNA-sequencing (snRNA-seq) and single-nucleus assay for transposase-accessible chromatin using sequencing (snATAC-seq). This comprehensive dataset comprises near two million nuclei from eight brain regions-prefrontal cortex (PFC), hippocampal formation, striatum, thalamus, hypothalamus, midbrain, pons and medulla, and cerebellum-of 13 cynomolgus monkeys (*Macaca fascicularis*), spanning the entire adult lifespan including exceptionally old individuals up to 29-years old. Through this atlas, we have investigated the molecular mechanisms underlying general and region-specific alterations in cell type composition, cell state and multicellular interactions across multiple stages of adulthood in these primates. Our analysis identified potential pathways and transcription factors (TFs) linked to brain aging and longevity, providing valuable insights into the cellular and molecular mechanisms underlying aging-related cognitive decline and neurodegeneration. We have also explored how aging-related modifications in chromatin accessibility within monkey brain cells correlate with human genetic risk variants implicated in cognition and neurodegeneration. Besides, by conducting a meta-analysis of publicly available human snRNA-seq datasets, we were able to highlight both the similarities and distinctions in transcriptional profiles between primate brain aging and AD.

The NHPABC serves as a foundational resource for generating novel hypotheses on primate brain aging, potentially supporting the development of therapeutic strategies aimed at improving brain health in older primates, including humans. This atlas is accessible via an interactive online portal at https://db.cngb.org/stomics/nhpabc/, facilitating exploration and integration with other datasets.

## RESULTS

### Building a NHP brain cell atlas across the adult lifespan

We selected the cynomolgus monkey because of its widespread use in biomedical research^14,17^. We dissected eight brain regions from 13 female euthanized monkeys, representing four age stages: ‘young adult’ (5-6 years old, *n* = 4), ‘middle-aged’ (10-12 years old, *n* = 3), ‘old’ (22-23 years old, *n* = 3), and ‘exceptionally old’ (28-29 years old, *n* = 3). The ‘exceptionally old’ group represents the maximum lifespan observed in cynomolgus monkey colonies in captivity and is quite rare^18^. The eight regions included the telencephalon (PFC, hippocampal formation, and striatum), diencephalon (thalamus and hypothalamus), mesencephalon (midbrain) and rhombencephalon (pons and medulla, and cerebellum) (**Figure 1A and Table S2**).

**Figure 1.**
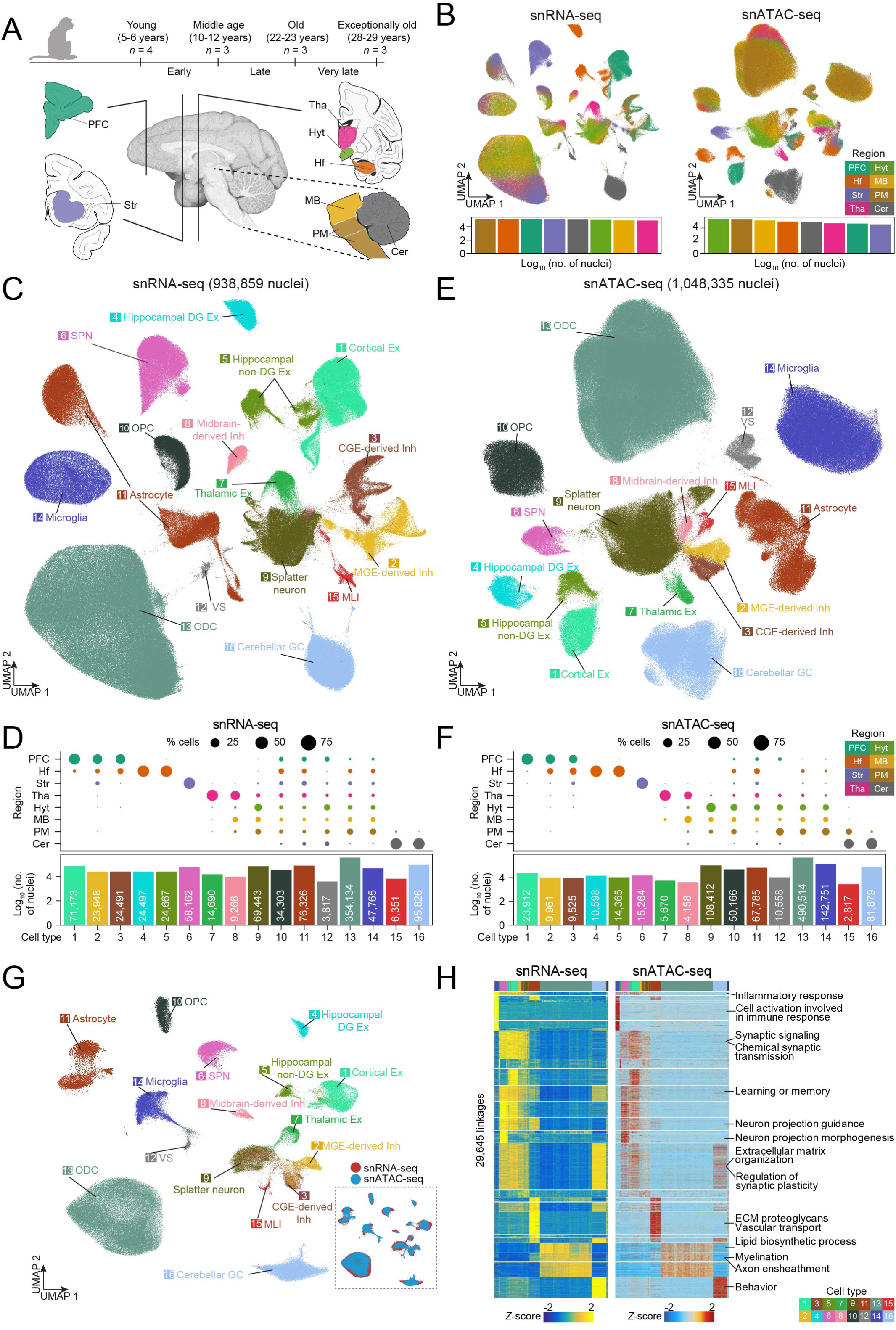
Generation of a cell atlas across eight regions of the aging monkey brain. (A) Schematic representation of the brain regions and the monkey age groups analyzed in this study. Top: the study was conducted on a cohort of 13 female monkeys divided into four age groups: young (5-6 years, *n*=4), middle age (10-12 years, *n*=3), old (22-23 years, *n*=3) and exceptionally old (28-29 years, *n*=3). Bottom: sagittal and coronal views of the monkey brain illustrating the specific regions analyzed. (B) UMAP visualization of all nuclei profiled by snRNA-seq (left) or snATAC-seq (right). The bar plots below indicate the number of nuclei profiled for each region. Dots and bars are coloured by region as indicated in the legends on the right-hand side. (C) UMAP visualization of 938,859 snRNA-seq profiles delineating 16 main brain cell populations. Dots are coloured by cell type. (D) Top: bubble plot showing the distribution of the main cell types identified in each region for snRNA-seq; the dot size indicates the percentage of cells, and the colour represents the region. Bottom: bar plot showing the number of cells identified for each annotated cluster in snRNA-seq; the colour represents the cell identity and the numeric labels correspond to the clusters shown in (C). (E) As in (C), UMAP visualization of 1,048,335 snATAC-seq profiles. (F) As in panel (**D**), for snATAC-seq. (G) UMAP visualization of the integrated snRNA-seq and snATAC-seq datasets, with each dataset downsampled to 100,000 cells. Dots are coloured by cell type (top) or omics modality (bottom right). (H) Heatmap showing the DEGs (left) and differentially accessible cCREs (right) from the integrated snRNA-seq and snATAC-seq data. Representative gene categories identified by functional enrichment are indicated on the right-hand side. Abbreviations: PFC, prefrontal cortex; Hf, hippocampal formation; Str, striatum; Tha, thalamus; Hyt, hypothalamus; MB, midbrain; PM, pons and medulla; Cer, cerebellum; Ex, excitatory neuron; Inh, inhibitory neuron; MGE, medial ganglionic eminence; CGE, caudal ganglionic eminence; DG, dentate gyrus; SPN, spiny projection neuron; OPC, oligodendrocyte progenitor cell; VS, vascular and stromal cell; ODC, oligodendrocyte; MLI, molecular layer interneuron; GC, granule cell.

All tissues were profiled by snRNA-seq and snATAC-seq with the DNBelab C4 droplet-based single-cell omics platform^14^. After quality control, we obtained 938,859 nuclei for snRNA-seq, varying from 82,817 in the thalamus to 146,473 in the pons and medulla, and 1,046,458 nuclei for snATAC-seq, varying from 38,299 in the striatum to 293,247 in the hypothalamus. On average, we detected 1,538 genes and 3,137 unique molecular identifiers (UMIs) per nucleus in the snRNA-seq, and 13,185 DNA fragments derived from transcription start sites (TSS) at 10.8 per nucleus in the snATAC-seq. Integrating the snRNA-seq information from all brain regions, we identified 16 main cell clusters in the global visualization of cell clustering using uniform manifold approximation and projection (UMAP) (**Figure 1B-D, S1A and Table S2**). Among the neuronal clusters, we noticed a group of neurons displaying both excitatory (Ex) and inhibitory (Inh) signatures mostly in non-telencephalic regions, referred to as splatter neurons^3^, enriched in the thalamus, hypothalamus, midbrain and pons and medulla. In addition, we identified all the expected non-neuronal cell types (astrocytes, microglia, oligodendrocytes, oligodendrocyte progenitor cells [OPCs] and vascular and stromal cells). Analysis of the snATAC-seq data revealed the same 16 main cell types (**Figure 1B, E, F, S1B and Table S2**). Cell composition in each region was consistent across ages and individuals in both datasets (**Figure S1C**), indicating a robust sampling and processing procedure. We also integrated the transcriptomic and epigenomic data resulting in overlapping cell clusters (**Figure 1G**). Moreover, the correlation of candidate cis-regulatory elements (cCREs; *P* < 0.01) in the main cell types with nearby expressed genes demonstrated the expected cell type correspondences across the two datasets (**Figure 1H**).

Hence, we have generated a large multi-region dataset for systematically characterizing the cell types composing the cynomolgus monkey brain at various stages of the adult lifespan until an exceptionally old age.

### A cell census of the cynomolgus monkey brain

To perform a high-resolution cell subtype classification, we integrated and reclustered the individual region snRNA-seq and snATAC-seq datasets. The regional cell subtype identities were determined by assessing gene expression and locus chromatin accessibility of canonical markers^3,6^ (*see website*).

This led to the identification of 89 neuronal subtypes in the snRNA-seq and a moderately smaller number in the snATAC-seq (**Figure 2A, B, S2A and Table S3**; *see website for a description of selected neuronal subtypes*). Among other things, our analysis of the splatter neurons from all non-telencephalic regions showed 29 neuronal populations related to specialized functions including multiple peptidergic hypothalamic neurons and an undefined cluster (**Figure 2B**; *see website*). Coronal hemibrain sections of a young adult cynomolgus monkey analyzed by Stereo-seq^17,19^ validated the observed distribution of selected neuronal subtypes identified in the snRNA-seq (**Figure S2B**).

**Figure 2.**
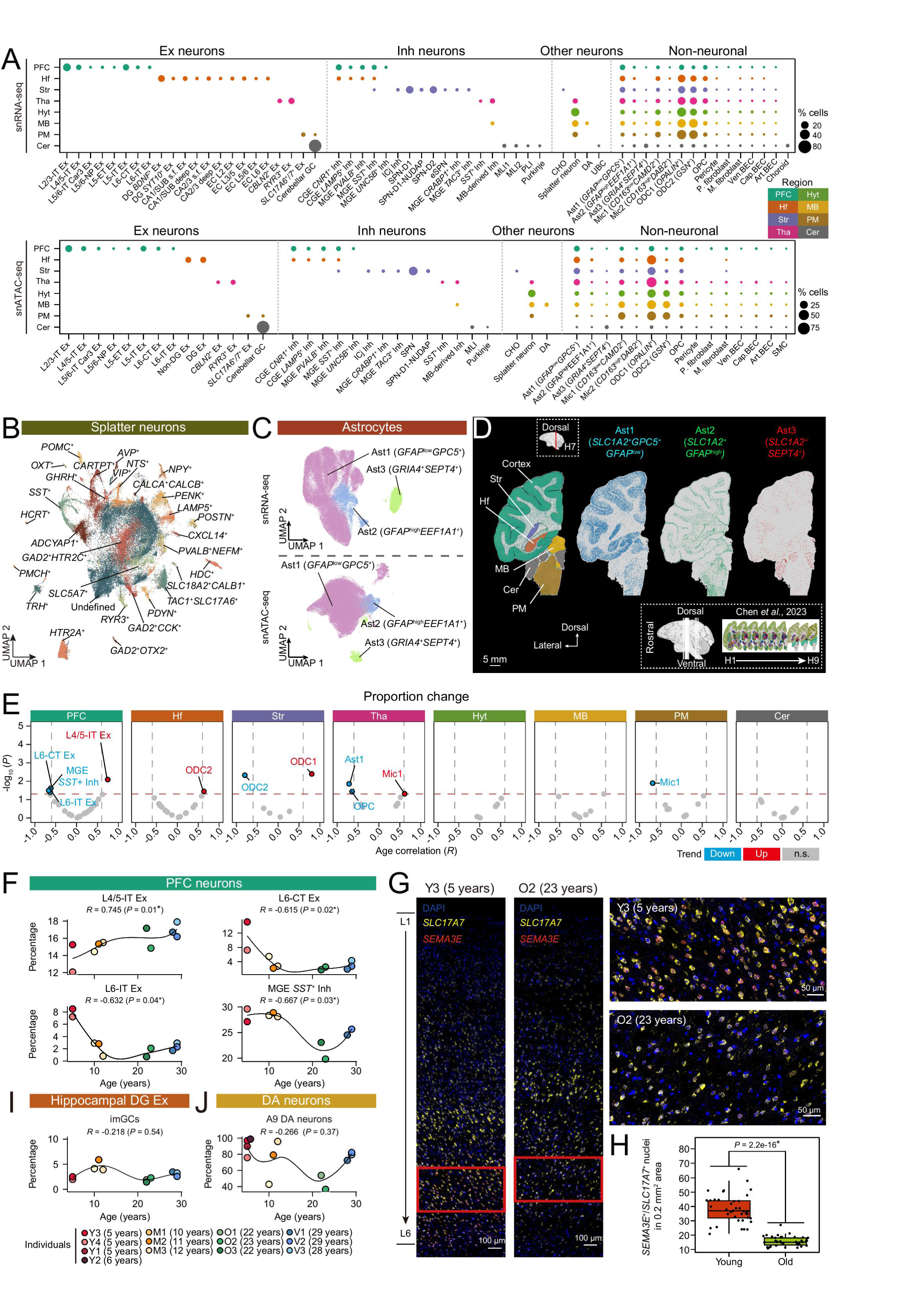
High-resolution spatiotemporal cell landscape of the aging monkey brain. (A) Bubble plot showing the distribution of each cell subtype (except for splatter neurons) across the eight brain regions analyzed by snRNA-seq (top panel) and snATAC-seq (bottom panel). The dot size indicates the percentage of cells, and the colour represents the region. (B) UMAP visualization of the reclustered splatter neurons from thalamus, hypothalamus, midbrain, and pons and medulla. The representative marker genes used to identify each population are indicated in the plot. (C) UMAP visualization of astrocyte subtypes from all brain regions in snRNA-seq (top) and snATAC-seq (bottom). The representative marker genes used to identify each population are indicated in the plot. (D) Spatial visualization of astrocyte subtypes (represented by the indicated marker genes) projected onto a hemi-brain section (section H7) from a five-year-old monkey using Stereo-seq data taken from Chen *et al.*. The schematic at the bottom right shows the anatomical position of the analyzed slides. Scale bar, 5 mm. (E) Volcano plots showing the correlation between the proportions of the indicated cell subtypes and the chronological age. The cell subtypes displaying age-related changes are indicated in each plot, with the blue label indicating the downregulated and the red indicating the upregulated ones. Those coloured in gray indicate no significant difference. *Q* values were calculated using a permutation test. (F) Line charts showing the correlation between the proportions of the PFC neuronal subtypes and the chronological age. The dots were coloured by individual. *P* value was calculated using a permutation test. (G) Representative images from smFISH detection of *SLC17A7* (yellow, for pan-Ex neurons) and *SEMA3E* (red, for L6 Ex neurons) in the PFC of young (individual Y3, 5 years-old) and old (individual O2, 23 years-old) monkeys. The right panels represent the magnification of the areas indicated by the red boxes in the left panels. Scale bars, 100 μm (left) and 50 μm (right). (H) Box plot comparing the numbers of *SLC17A7*^+^/*SEMA3E*^+^ L6 Ex neurons in the PFC from young and old animals (*n*=2 individuals from each age stage). The dots indicate the areas included in the analysis. (I) Line chart showing the correlation between the proportion of hippocampal DG Ex neurons and chronological age. The dots were coloured by individual. *P* value was calculated using a permutation test. (J) As in panel (**I**), for DA neurons. Abbreviations: Art, arterial; Ast, astrocyte, BEC, brain endothelial cell; CA, cornu ammonis; Car, claustrum-like; Cap, capillary; CHO, Cholinergic neuron; Choroid, choroid plexus; CT, corticothalamic; DA, dopaminergic neuron; EC, entorhinal cortex; ET, extratelencephalic; ICj, islands of Calleja; imGC, immature granule cell; IT, intratelencephalic; M. fibroblast, meningeal fibroblast; Mic, microglia; NP, near-projecting; P. fibroblast, perivascular fibroblast; PLI, Purkinje layer interneuron; SMC, smooth muscle cell; SUB, subiculum; UBC, unipolar brush cell; Ven, venous.

By integrating the non-neuronal cell types across brain regions, we found three types of astrocytes, two types of microglia, two types of ODCs and six types of vascular and stromal cells, but there was no clear subdivision of OPCs (**Figure 2C and S2C**). Among the astrocytes, we detected a major Ast1 (*GFAP*^low^ *GPC5*^+^) population, and two smaller clusters, Ast2 (*GFAP*^high^ *EEF1A1*^+^) and Ast3 (*GRIA4*^+^ *SEPT4*^+^). Stereo-seq showed widespread brain distribution of Ast1, preferential distribution of Ast2 in the white matter and L1-pial border of the cortex, and enrichment of Ast3 in the cerebellum (**Figure 2D**). The distribution of Ast1 and Ast2 resembles that of the previously described ‘protoplasmic’ and ‘interlaminar’ or ‘fibrous’ astrocytes^20^. Microglia comprised quiescent microglia (Mic1; *CD163*^low^*CAMD2*^+^) and a smaller cluster with an activated macrophage signature (Mic2; *CD163*^high^ *DAB2*^+^) (**Figure S2C**). Both subtypes showed a brain-wide distribution in the snRNA-seq and when analyzed by Stereo-seq (**Figure S2D**). ODCs could be subdivided into *OPALIN^+^* ODC1, responsible for myelin formation, and a *GSN^+^* ODC2, which are involved in regulating cell motility during myelin wrapping^3^ **(Figure S2C**). In the Stereo-seq sections, ODC1 were largely enriched in the white matter, while ODC2 were more broadly distributed (**Figure S2D**). Vascular and stromal cells included pericytes, perivascular fibroblasts, and meningeal fibroblasts, as well as venous, capillary, and arterial brain endothelial cells (**Figure S2C**). A small population of smooth muscle cells was only captured in the snATAC-seq.

This cellular landscape, integrated with prior knowledge from studies on mice and humans, provides a valuable resource for investigating the effects of aging on specific cell types and brain regions in the cynomolgus monkey.

### Aging-associated changes in cell populations

Neurons, as post-mitotic cells with long lifespan, and glial cells, experience various stresses throughout life. This makes them vulnerable to cell death, especially in the context of neurodegeneration^4,12,13^. However, the question of whether specific cell types are more susceptible to death during aging has not been systematically examined across multiple brain regions in the same NHP individuals. Utilizing our multi-region dataset of cynomolgus monkeys, we sought to address this question. To avoid potential biases in the calculations of cell proportions due to individual variation or sampling, we applied a permutation test (*R* > 0.6 or *R* < –0.6, *Q* < 0.05 and number of cells > 10 in each sample).

Among PFC Ex neurons, there was a selective age-related decline in the proportion of layer 6 (L6)-IT (intratelencephalic) and L6-CT (corticothalamic) Ex neurons especially in the transition from young to middle-aged and old monkeys (**Figure 2E, F**). In contrast, the proportion of layer 4/5 (L4/5)-IT Ex neurons increased. The decrease in L6-CT Ex neurons was validated using single-molecule fluorescence *in situ* hybridization (smFISH) for *SEMA3E* and *SLC17A7* co-staining (**Figure 2G, H**). Among PFC Inh neurons only medial ganglionic eminence (MGE)-derived *SST*^+^ neurons displayed a decline in old monkeys (**Figure 2E, F**), as reported in the dorsolateral PFC of older humans and AD patients^4,13^. We did not find a substantial decline of neurogenesis with aging in the hippocampus, as indicated by the maintenance of the pool of immature granule cells (imGC, *DCX*^+^ *PROX1*^+^)^21,22^ (**Figure 2I**; *see website*). This is consistent with any reduction in these cells being marginal after reaching the adult stage. However, we observed a reduction of A9 dopaminergic (DA) neurons (*TH^+^ SOX6*^+^) In the midbrain of old monkeys, in line with the increased prevalence of Parkinson’s disease in older humans^23^ (**Figure 2J**; *see website*). Notably, there was a smaller reduction of A9 DA neurons and MGE-derived *SST*^+^ Inh neurons from the PFC of exceptionally old monkeys. This suggests that the age-related changes in these individuals follow a distinct trajectory.

Most non-neuronal cell types did not exhibit any notable trend, but we observed specific ones with variable behavior (**Figure 2E**). Mic1 increased with aging in the thalamus but decreased in the pons and medulla. ODC1 and 2 increased and decreased, respectively, in the striatum, whereas ODC2 increased in the hippocampal formation. Furthermore, OPCs and Ast1 decreased selectively in the thalamus.

Therefore, alterations in the proportion of cell populations in the aging cynomolgus monkey brain appear to be rather selective. Additionally, we observed a recurring theme throughout the following sections: the partial preservation of certain molecular features affected by aging in elite survivors at exceptionally old ages.

### Aging-associated gene expression dynamics in neurons

We then focused on deciphering the spatiotemporal transcriptomic logic of different neuronal subtypes across the cynomolgus monkey brain. To define aging-associated differentially expressed genes (DEGs) in neurons we used two approaches: i) a chronological linear model to identify progressive aging-related DEGs (pDEGs); 2) a pairwise comparison between adjacent age groups for detecting stage-specific DEGs (sDEGs). The motivation for this was to gain a deeper understanding of the kinetics of changes and to explore potential distinct molecular trajectories in the exceptionally old group. We detected 5,319 pDEGs (age coefficient > 0.005 or < –0.005^24,25^). Among the different comparison groups, we observed the following numbers of sDEGs (log_2_[fold change] > 0.25): young adult to middle-aged (‘early stage’) 4,246, middle-aged to old (‘late stage’) 4,229 and old to exceptionally old group (‘very late stage’) 2,723 (**Figure 3A**; *see website*). DEGs in both analyses were not biased by differences in the number of captured cells or detected genes (**Figure S3A**). The neuronal subtype with the largest number of pDEGs and sDEGs was *SLC17A6*/*7*^+^ Ex neurons in the pons and medulla, suggesting an enhanced vulnerability of this region in aging. Most sDEGs were stage-specific, with the largest proportion at the early stage (**Figure S3B, C**). Moreover, as in previous reports of mouse tissue aging^24^, there were more downregulated than upregulated pDEGs and sDEGs (**Figure 3A and S3D**). A notable exception were sDEGs at the very late stage, which displayed the opposite trend. The downregulated pDEGs with the highest fold change were *NPY* and *SST* in *SST*^+^ Inh neurons from the thalamus (**Figure S3E**). Functional enrichment analysis showed that upregulated sDEGs at the very late stage encompassed categories associated with endocytosis, autophagy, apoptosis, protein folding, cellular respiration, and neurotransmitter secretion (**Figure S3F**). This is particularly relevant because neurodegeneration is associated with impairment of these processes^26^.

**Figure 3.**
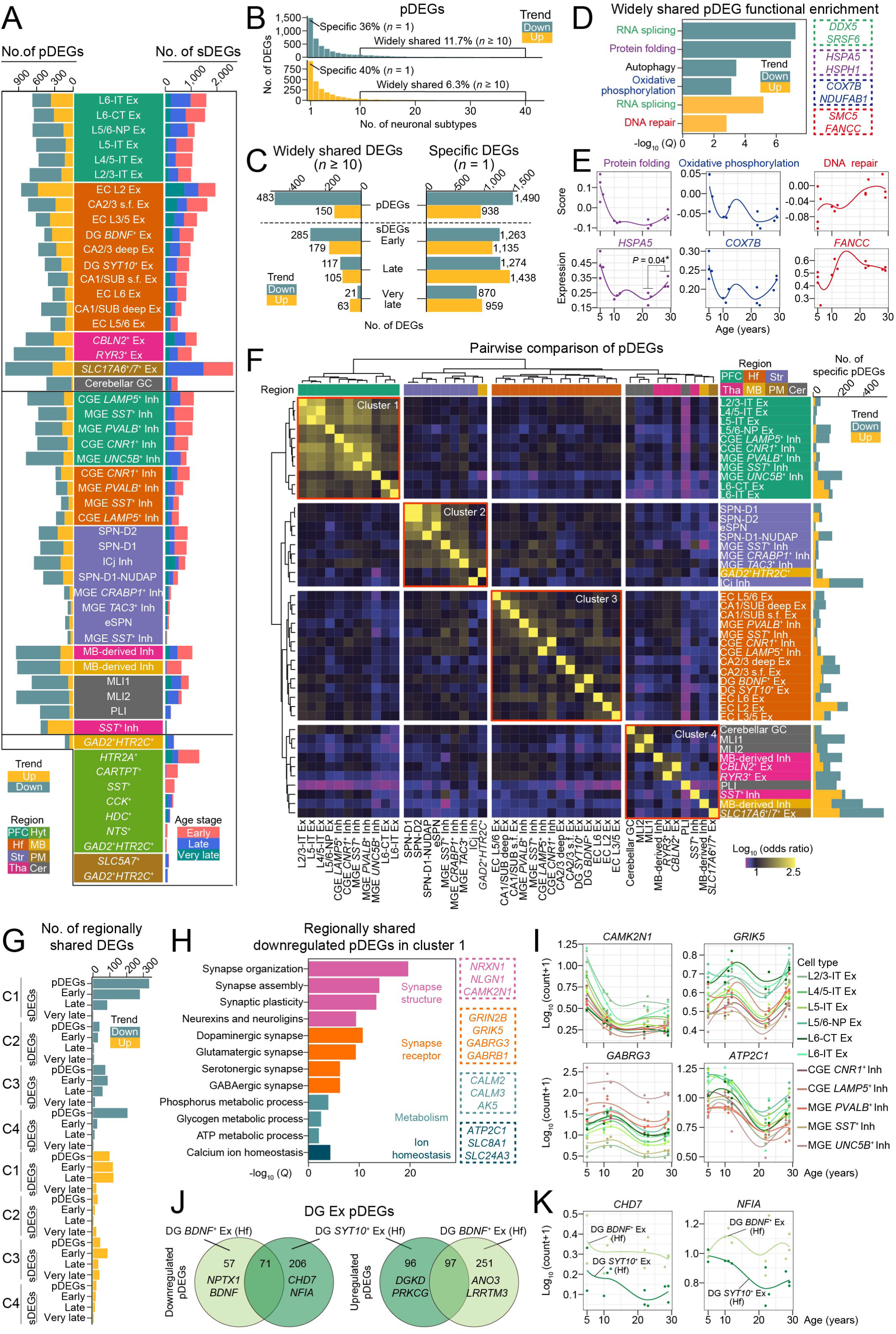
Analysis of aging-associated gene expression dynamics in neurons. (A) Stacked bar plot showing the number of pDEGs (left) in 43 neuronal subtypes and sDEGs (right) in 53 neuronal subtypes. The central bar indicates the cell subtype and is coloured by region. Bar plots on the left and right are coloured by the pDEG expression trend (upregulated or downregulated) and by each age-stage comparison of sDEGs (early, young/middle age; late, middle age/old; very late, old/exceptionally old), respectively. (B) Histogram showing the frequency distribution of downregulated (top panel) or upregulated (bottom panel) pDEGs across different cell subtypes, coloured by the expression trend. The plot includes the proportion of genes that are widely shared (identified in at least 10 cell subtypes) or cell subtype-specific (identified in only one cell subtype), for both the downregulated and upregulated pDEGs. (C) Bar plots showing the number of widely shared (left) and cell-type specific (right) DEGs, coloured by the expression trend. (D) Bar plot showing the functional enrichment results of widely shared DEGs. The dotted box on the right contains the representative genes that correspond to the functional terms or pathways highlighted in the same colour in the heatmap on the left. The colour scale represents the significance (−log_10_[*Q*]) of the enriched terms for downregulated (green) and upregulated (yellow) genes with aging. (E) Line charts showing the module score of the indicated pathways (top panels) and the expression levels (bottom panels) of representative genes related to each pathway in all the neuronal cells from each individual. The dots represent each individual. (F) Hierarchical heatmap showing the clustering of all neuronal populations based on pDEG similarity. The level of similarity (−log_10_ [odds ratio]) is depicted by the colour scale. Region identity is shown on the right-hand side legend. The bar plot on the right shows the number of cell-type specific pDEGs in the corresponding neuronal subtype and is coloured by expression trend. (G) Bar plot showing the numbers of regionally shared pDEGs and sDEGs in each cluster indicated in panel (**F**). (H) Bar plot showing the functional enrichment results of regionally shared pDEGs in cluster 1. The dotted boxes on the right contain specific genes corresponding to the functional terms highlighted in the same colour. (I) Line charts showing the expression levels of genes related to synapse structure (*CAMK2N1*), synapse receptor (*GRIK5*, *GABRG3*) and ion homeostasis (*ATP2C1*) in all PFC neuronal subtypes from each individual. The dots represent each individual. (J) Venn diagram showing the overlap between the upregulated and downregulated early pDEGs of hippocampal DG *BDNF*^+^ Ex and *SYT10*^+^ Ex neurons. The numbers of overlapping and non-overlapping genes are labelled on the plot. (K) Line charts showing the expression levels of *CHD7* and *NFIA* at different ages in hippocampal DG *SYT10*^+^ Ex neurons. The dots represent each individual.

We next examined the specificity of the DEGs by performing two extra analyses. First, we searched for DEGs shared among neuronal subtypes in any of the eight brain regions. The vast majority of pDEGs and sDEGs were exclusive to only one or few neuronal subtypes, whereas a small proportion (‘widely shared’) appeared in ≥10 subtypes (**Figure 3B, C and S3G**). The downregulated widely shared pDEGs, belonged to pathways related to protein folding, autophagy, and oxidative phosphorylation (**Figure 3D, E**). Multiple downregulated pDEGs belonging to these categories (*HSPA5*, *COX7B*) were partially reversed in exceptionally old monkeys, substantially intersecting with the above-mentioned upregulated sDEGs at the very late stage (**Figure S3H**). Upregulated widely shared pDEGs included genes related to DNA repair. In addition, we observed pDEGs belonging to RNA splicing (*HNRNPA2B1*, *SRSF7*) with variable dynamics across different brain regions but substantially upregulated in the PFC and the pons and medulla (**Figure S3I, J**). Second, we did a pairwise comparison of pDEGs based on the odds ratio of overlap between each neuronal subtype across all regions (**Figure 3F and S3K**). This showed four clusters ranking from higher to lower similarity. Cluster 1 were PFC neurons, cluster 2 mostly corresponded to striatal neurons, cluster 3 were hippocampal formation neurons, and cluster 4 consisted of heterogeneous neuronal populations. Similar conclusions were obtained with sDEGs (**Figure S3L**). We also noticed that L6-CT Ex neurons displayed a larger number of sDEGs at the early stage than other PFC neuronal subtypes. The most significantly enriched category for upregulated sDEGs in these neurons was developmental-related genes (**Figure S4A**), suggesting that they are either still maturing or dedifferentiating.

We searched the ‘regionally shared’ DEGs, defined as the pDEGs and sDEGs appearing in more than half of the cell subtypes in each of the clusters above. As expected, the pDEGs in cluster 1 were the largest group. Downregulated pDEGs in this cluster were more abundant than upregulated ones and were enriched in pathways related to synaptic organization and function, metabolism, and Ca^2+^ homeostasis (**Figure 3G-I and S4B**). For instance, *CAMK2N1*, an inhibitor of calcium/calmodulin-dependent protein kinase II (CaMKII) and the Ca^2+^ channel *ATP2C1* were downregulated at the late stage in most PFC neuronal subtypes. CaMKII is a key synaptic signaling molecule for learning and memory^27^. Multiple glutamate receptors (*GRIK5, GRIN2B*) in PFC Ex neurons and GABA receptors (*GABRA1*, *GABRG3*) in PFC Inh neurons were also downregulated at the late stage. Notably, *CAMK2N1* was consistently downregulated in neurons from the hippocampal formation but glutamate and GABA receptors were not (**Figure S4C**). We also compared the regionally shared pDEGs in cluster 1 with the widely shared pDEGs, observing an ∼55% overlap (**Figure S4D**). This indicates that among all the profiled neuronal subtypes in the monkey aging brain those in the PFC are the major contributors to widely shared DEGs.

In addition, we studied the changes in specific paired or individual neuronal subtypes. We detected a higher number of downregulated DEGs in *SYT10*^+^ Ex neurons from the hippocampal dentate gyrus (DG) compared to their analogous *BDNF*^+^ subtype, especially at the late stage (**Figure 3J, K**). For example, *CHD7* (chromodomain helicase DNA binding protein 7), which is associated with the maintenance of hippocampal neural progenitor cell quiescence in rodents^28^, decreased only in *SYT10*^+^ Ex neurons. *NFIA* (nuclear factor IA), which is downregulated in human hippocampal imGCs upon aging^21^, followed a similar pattern. While midbrain-derived Inh neurons in the thalamus and midbrain presented shared DEGs such as the upregulated *SLC6A15*, which is associated with increased anxiety-like behavior in mice^29^, aging was also associated with distinct changes (**Figure S4E**). For example, the receptor tyrosine kinase *KIT*, which regulates axonal extension in development^30^, was selectively reduced in the thalamus. Moreover, the GABA receptors *GABRA1* and *GABRB2* were selectively decreased in Inh neurons from the islands of Calleja (ICj) but no other neurons from the striatum (**Figure S4F**). The ICj are one of the main areas processing the brain reward pathway, whose modulation is altered in older humans^31^. Further, changes in Purkinje layer interneurons (PLI) were distinctive compared to other cerebellar neurons. Among the specific pDEGs, we identified a decline in genes related to synaptic transmission including *PSEN1* (presenilin 1); presenilins are essential for regulating neurotransmitter release^32^. We also noticed a reduced *LARGE1* expression, which has been previously implicated in inherited cerebellum dysplasia (dystroglycanopathy)^33^, at the late and very late stages. Likewise, downregulated pDEGs in *SLC17A6*^+^/*7*^+^ Ex neurons from the pons and medulla were enriched in regulators of trans-synaptic signaling (*PTPRS*, *CACNB4*). This is consistent with the critical role of this region integrating signals from other brain areas and the spinal cord. Likewise, we observed enrichment in downregulated genes related to axon guidance (*ROBO1*, *NRCAM*). For hypothalamic peptidergic neurons, we noticed a consistent downregulation between individuals of the neuropeptides *ADCYAP1* (adenylate cyclase activating polypeptide 1) and *GHRH* (growth hormone-releasing hormone with aging) in the corresponding neuronal subtypes (**Figure S4G**).

In summary, the transcriptional changes induced by aging in the cynomolgus monkey brain are predominantly specific to neuronal subtypes. However, there are also region-specific effects that impact multiple neuronal subtypes simultaneously, particularly in the telencephalon.

### Aging-associated gene expression dynamics in non-neuronal cells

We applied the same spatiotemporal analyses to non-neuronal cells. Since the numbers of nuclei were more limited, we pooled the different subtypes of vascular and stromal cells in all regions and excluded Mic2 regional subtypes for the calculations. Altogether, we identified 4,994 global pDEGs and 7,075 global sDEGs across the eight brain regions, with no bias due to differences in the number of captured cells or detected genes (**Figure 4A and S5A***; see website*). DEGs were more abundant in non-telencephalic glial subtypes. Most sDEGs were stage-specific, and there were also more downregulated than upregulated DEGs except for sDEGs at the very late stage (**Figure S5B**). The downregulated and upregulated pDEGs with the highest fold change were *COL21A1* in PFC Ast2 and *SPARCL1*^34^ in midbrain Ast1, respectively (**Figure S5C**). We then searched for pDEGs and sDEGs shared between different regional glia subtypes. The shared DEGs (appearing in more than half of all regional subtypes in each main glial cell type) only accounted for a small proportion of DEGs (**Figure S5D**). Pairwise comparison of pDEGs across all glial subtypes revealed strong cell subtype specificity, with greater similarity observed between ODCs from different regions compared to other glial cells (**Figure S5E, F**).

**Figure 4.**
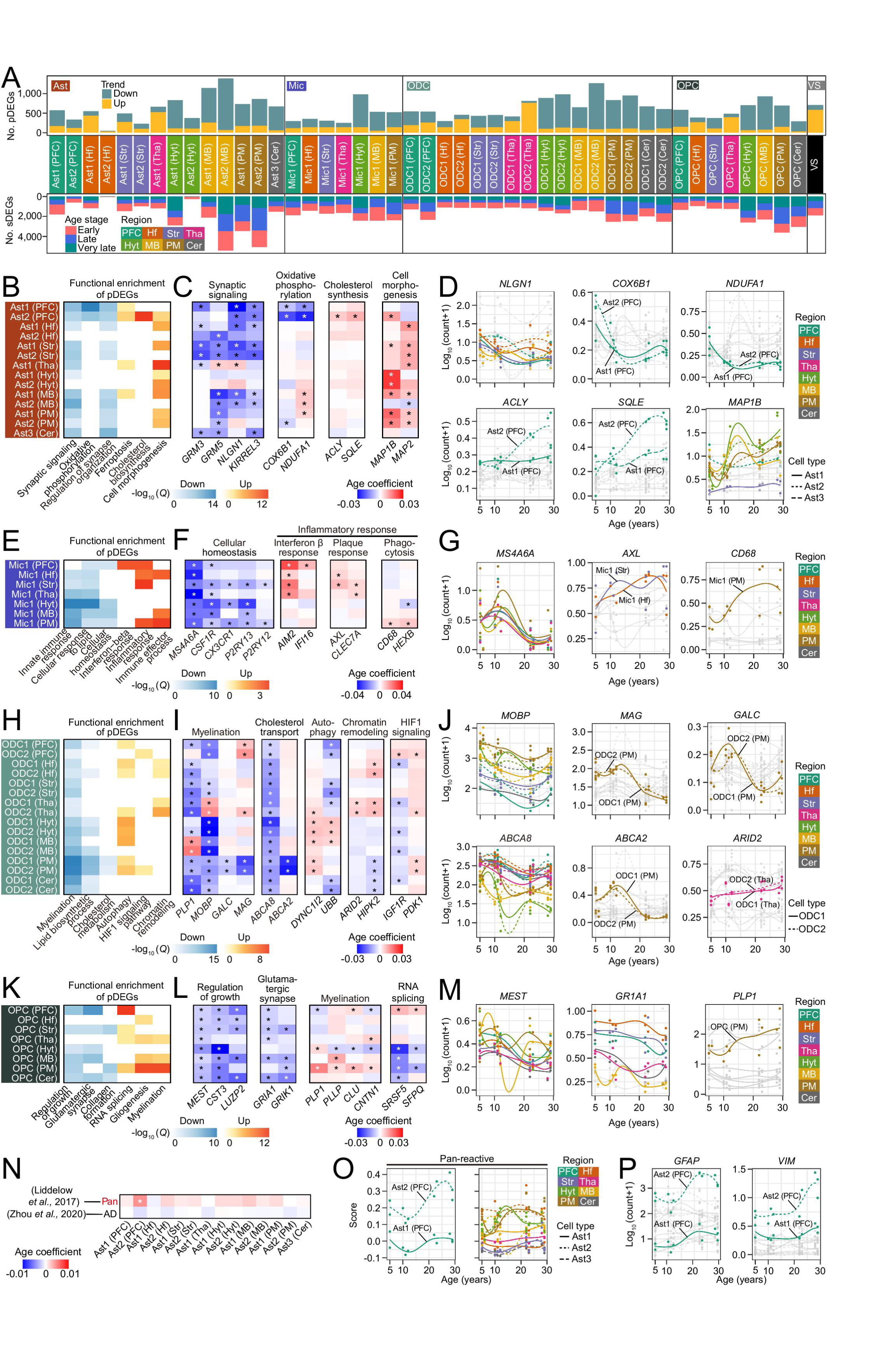
Analysis of aging-associated gene expression dynamics in non-neuronal cells. (A) Stacked bar plot as in Figure 3A, but for 46 non-neuronal cell subtypes. (B) Heatmap showing the functional enrichment results of pDEGs in astrocyte subtypes. The colour scale represents the significance (−log_10_[Q]) of the enriched terms for downregulated (blue) and upregulated (red) DEGs with aging. (C) Heatmap showing the age-coefficient of genes related to synaptic signaling, oxidative phosphorylation, cholesterol synthesis and cell morphogenesis in astrocyte subtypes. The colour scale represents the age-coefficient. (D) Line charts showing the expression levels of genes related to synaptic signaling (*NLGN1*), oxidative phosphorylation (*COX6B1, NDUFA1*), cholesterol synthesis (*ACLY*, *SQLE*), and cell morphogenesis (*MAP1B*) across astrocyte subtypes from each individual. The dots represent each individual. Dots and lines are coloured by region, with each line representing a specific cell subtype. Dots and lines coloured in grey indicate a non-statistically significant trend. (E-G) As in Figure 4B-D, for microglia. (H-J) As in Figure 4B-D, for ODCs. (K-M) As in Figure 4B-D, for OPCs. (N) Heatmap showing the age-coefficient value for module scores of different reactive astrocyte states (PAN, AD) in astrocyte subtypes. The gene lists are taken from previous studies (*see website*). (O) Line charts showing the module score of genes related to the pan-reactive astrocyte state in astrocyte subtypes from each individual. Dots and lines are coloured by region, with each line representing a specific cell subtype. Dots represent each individual. (P) Line charts showing the expression levels of two pan-reactive astrocyte genes (*GFAP*, *VIM*) across astrocyte subtypes from each individual. The dots represent each individual.

Astrocytes provide support for neurons, participate in synapse regulation, and modulate axon myelination via crosstalk with OPCs and ODCs^35^. Functional enrichment analysis for pDEGs revealed a broad aging-related downregulation of genes associated with synaptic signaling in Ast1 and Ast2 from different regions. This included glutamatergic signaling receptors (*GRID2*, *GRM3*/*5*) and genes related to synapse organization (*NLGN1*, *KIRREL3*) (**Figure 4B-D**). Likewise, genes associated with oxidative phosphorylation (*COX6B1, NUDFA1*) were downregulated with aging in PFC Ast1 and Ast2, whereas genes related to cholesterol biosynthesis (*ACLY*, *SQLE*) were upregulated in PFC Ast2. Excessive cholesterol synthesis in astrocytes carrying the *APOE4* allele has been proposed to contribute to AD pathogenesis^36^. In addition, pDEGs related to cell morphogenesis, including cytoskeletal components (*MAP1B*, *MAP2*), were broadly upregulated in Ast1 and Ast2.

Microglia clear brain debris (cells, myelin, and aggregates) through phagocytosis, prune synapses, and support neuronal survival and remyelination^37^. All regional Mic1 subtypes were enriched in downregulated pDEGs associated with cellular homeostasis (*MS4A6A*, *P2RY13*), particularly in the hypothalamus, midbrain, and pons and medulla (**Figure 4E-G**). Upregulated pDEGs were predominantly associated with inflammatory responses and, in contrast, displayed region-specific effects. For instance, interferon β response genes (*IFI16*, *AIM2*) were upregulated in the PFC, while amyloid plaque-associated genes (*AXL*, *CLEC7A*) were upregulated in the hippocampal formation, striatum, and thalamus. Interestingly, phagocytosis-related genes (*CD68*, *HEXB*) were mainly upregulated in the pons and medulla. ODCs generate the myelin sheaths that encircle the axonal bundles and ensure efficient electrical signal transmission^38^. Downregulated pDEGs in ODCs were enriched in genes related to myelination (*MOBP*, *PLP1*) and lipid biosynthetic process (*ABCA8*), indicative of an abnormal cell maturation state, with the former category being more broadly affected (**Figure 4H-J**). The pons and medulla displayed strong down-regulation of genes in both categories. For example, the myelin component *GALC* and *MAG*, as well as the cholesterol efflux transporter *ABCA2*, were more reduced in the pons and medulla at the late stage than in other regions. Aberrant lipid accumulation in ODCs has been described in AD patients^39^. Among the categories for upregulated pDEGs, HIF1 signaling (*IGF1R*, *PDK1*) was moderately upregulated in ODC2 from the PFC and pons and medulla. Hypoxia resulting from a compromised vasculature induces HIF1 signaling and delays ODC maturation^40^. We also noticed a variable behavior of genes related to autophagy and chromatin remodeling in different regions.

OPCs respond to demyelination by proliferating, migrating and subsequently differentiating into ODCs. Among the downregulated pDEGs in OPCs we found broad enrichment in categories associated with OPC homeostasis such as regulation of growth and glutamatergic synapse (*GRIA1*, *GRIK1*). Synaptic activity between neurons and OPCs modulates their differentiation into ODCs^41^. For upregulated pDEGs, categories associated with OPC differentiation (gliogenesis and myelination: *PLP1*, *CNTN1*) increased more substantially in the pons and medulla. The latter indicates a more active exit from the quiescent state compared to other regions (**Figure 4K-M**), consistent with the changes in ODCs. In addition, RNA splicing-related genes (*SRSF5*, *SFPQ*) were strongly upregulated in PFC OPCs and less obviously in the pons and medulla.

Changes in blood flow and the blood-brain barrier (BBB) are major contributors to brain aging and neurodegeneration^42,43^. Among the categories for downregulated pDEGs in vascular and stromal cells, we observed ribosomal subunits (*RPL5*, *RPS7*) and genes involved in extracellular matrix (ECM) organization (*BGN*, *LAMB2*) (**Figure S5G, H**). Upregulated pDEGs comprised genes belonging to membrane and transmembrane transport (*SLC15A5*, *SLC6A20*)^43^ and HIF1 signaling (*ALDOC*, *PRKCA*). We also noticed upregulation of the AD-associated genes *NLGN1* and *BNC2*. The former participates in developmental vasculogenesis^44^ while the latter regulates ECM organization and cytokine production in fibroblasts^45^.

In response to injury or disease, glial cells undergo molecular and functional remodeling, often referred to as disease-associated or reactive states^46–53^. To ascertain whether monkey brain aging results in similar cell state transitions, we searched for previously reported transcriptomic signatures of reactive glia in our dataset (**Figure 4N, O**; *see website*). We did not observe the activation of a pan-reactive^46^ or AD-associated phenotype^47^ in aging NHP astrocytes. One exception, however, were Ast2 in the PFC, which showed a pan-reactive signature at the late stage. The reactive state of this astrocyte subtype was further confirmed by the induction of *GFAP* and *VIM* (**Figure 4P**). This is consistent with previous observations in mice showing that PFC Ast2, which match the spatial distribution of PFC Ast2 in monkey, are more affected in aging than Ast1^8^. We did not observe either a clear transition of homeostatic microglia to disease-associated microglia (DAM: DA1, DA2 and IFN)^48^, neurodegenerative phenotype microglia (MGnD)^50^, or white matter-associated microglia (WAM)^51^. (**Figure S5I**). Similarly, there was no obvious induction of a disease-associated ODC (DAO) state, as defined in AD and multiple sclerosis (MS) patients^52^, or evidence of disease-associated ODCs (DOLs) described in an AD mouse model^53^.

We conclude that: i) similar to neurons, non-neuronal cells exhibit significant inter– and intra-regional heterogeneity in their responses to aging; ii) multiple lines of evidence suggest that myelination injury is more pronounced in the pons and medulla compared to other brain regions, which may explain the larger number of aging-related DEGs observed in neurons of these areas; iii) in contrast to human neurodegeneration, glial cells in the aging cynomolgus monkey brain do not show broad reactivity or polarization into prototypical phenotypes.

### Changes in regional multicellular crosstalk with aging

Currently open questions include how aging alters cellular communication in the primate brain, how this influences key cellular processes including synaptogenesis and myelination, and whether there are regional differences. To address these questions, we searched our cynomolgus monkey dataset for ligand-receptor interactions associated with the identified aging-associated DEGs (pDEGs and sDEGs). The interactions were determined by CellChat^54^. This identified 617 differentially expressed ligands and receptors corresponding to a total of 141,234 ligand-receptor interactions among the different cell types and regions (**Figure 5A**; *see website*). The largest number of aging-associated interactions were in the PFC and hippocampal formation, and the smallest was in the hypothalamus (**Figure S6A**). Moreover, telencephalic and non-telencephalic regions were more enriched in inter-neuronal and inter-glial interactions, respectively (**Figure S6B**). We determined the categories of cell-cell interactions altered by aging using functional enrichment analysis. Among them, we noticed synaptic signaling, growth factors, inflammatory response, ECM organization, and myelination with distinct enrichment in neurons or glia (**Figure 5B**).

**Figure 5.**
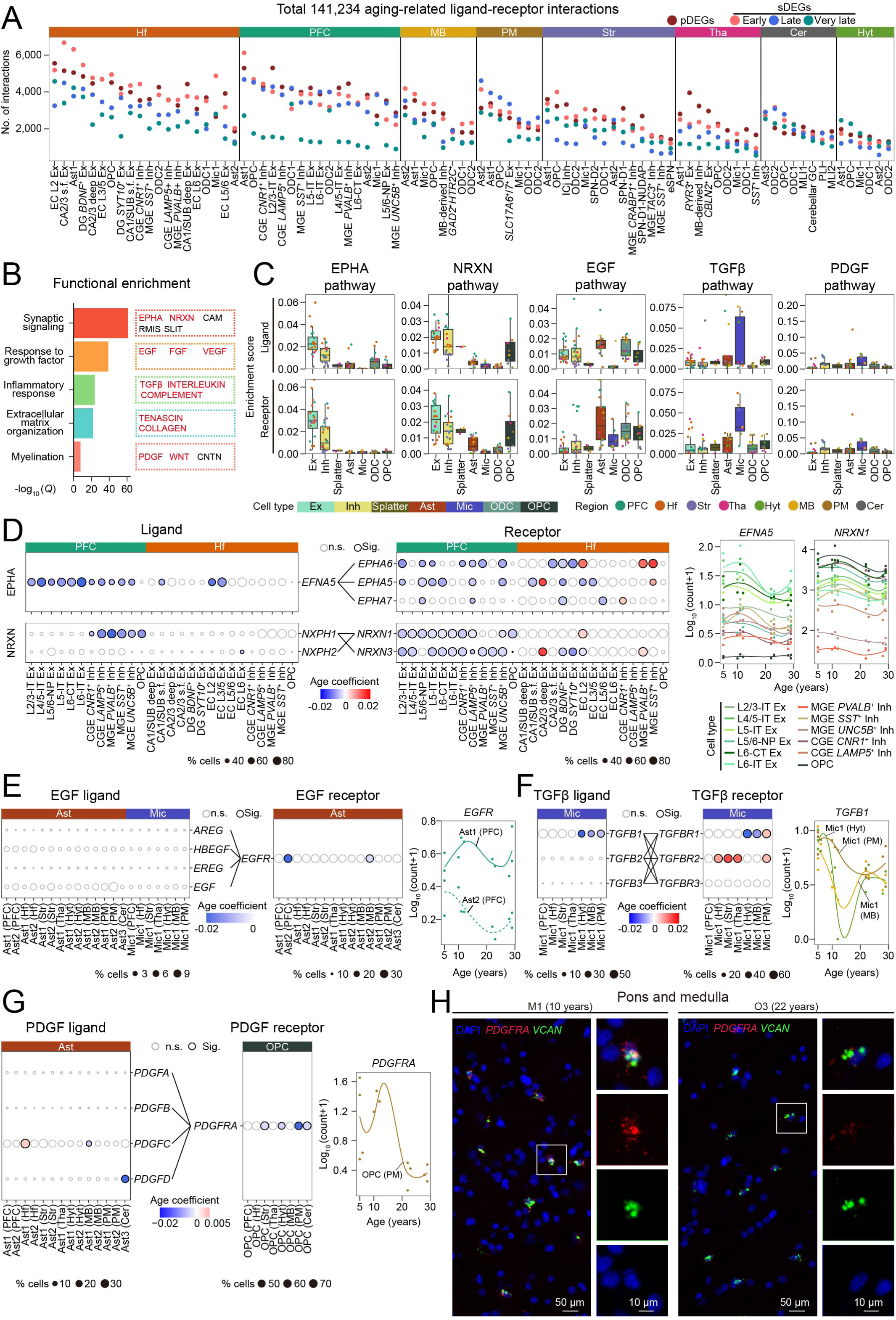
Brain-wide dynamic cell-cell communication changes with aging. (A) Dot plot showing the number of cell-cell interactions related to DEGs among the cell subtypes from each brain region. Region identity is indicated on the top. Dots are coloured by the DEG category. (B) Bar plot showing the functional enrichment results of 617 differentially expressed ligands and receptors. Specific genes related to the colour matched pathways are highlighted on the right-hand side. (C) Box plot showing the ligand (top row) or receptor (bottom row) enrichment score of specific pathways across different cell subtypes. The box plots are coloured by cell type, and the dots within the boxes by brain region. (D) Left: bubble plots showing the expression levels and age-coefficients for ligands (left) and receptors (right) in the EPHA and neurexin pathways in the indicated cell subtypes of PFC and hippocampal formation. Black lines connect ligands with their corresponding receptors. Right: line charts showing the expression levels of the indicated genes in the indicated cell subtypes. (E) As in Figure 5D, for EGF pathway in microglia and astrocytes. (F) As in Figure 5D, for TGFβ pathway in microglia. (G) As in Figure 5D, for PDGF pathway in astrocytes and OPCs. (H) Representative images from smFISH detection of *PDGFRA* (red) and *VCAN* (green) in the pons and medulla of middle-aged (individual M1, 10 years-old) and old (individual O3, 22 years-old) monkeys. The right panels are a magnification in the area indicated by the white box in the left panels. Scale bars, 50 μm (big view) or 10 μm (small view).

As part of synaptic signaling, we observed the downregulation of several pathways including ephrin and neurexin pathways (**Figure 5C, D and S6C**). Ephrins and their ligands are cell adhesion molecules involved in synaptogenesis^55^, whereas neurexins are synapse organizers^56^. The ephrin-A ligand *EFNA5,* its receptors *EPHA5*/*6*/*7*, the secreted neurexin ligands (neurexophilins) *NXPH1*/*2* and their receptors *NRXN1*/*3*, were mainly downregulated in Ex and Inh neurons from the PFC and to a lesser extent Ex neurons from the hippocampal formation. PFC OPCs also exhibited downregulation of the neurexin pathway. These effects were more prominent at the late and very late stage, consistent with the general downregulation of synapse-related genes in PFC neurons at those stages (see **Figure 3H** above). Growth factors play crucial roles in neuronal and glial homeostasis and their responses to stress or injury. *FGF12*/*14* were downregulated in PFC Ex and Inh neurons and *FGF13* only in PFC Inh neurons (**Figure S6C**). Loss of *Fgf13* is associated with axonal abnormalities leading to deficits in memory and learning in mice^57^. EGFR was selectively downregulated in PFC Ast2 at the late stage and partially reversed in exceptionally old monkeys (**Figure 5E**). EGF signaling triggers quiescent astrocytes into a reactive state in response to injury^58^. Although this contrasts with the enhanced reactivity in this astrocyte subtype with aging (see **Figure 4N** above), it may indicate a compensatory response to prolonged activation. Among the *VEGF* family, *VEGFA* was downregulated in astrocytes and OPCs from multiple regions (**Figure S6C**). The role of the VEGF family in neurodegeneration is complex, but growing evidence suggests that VEGFA provides protection against cognitive impairment in AD^59^.

Neurodegeneration is accompanied by an increased secretion of pro-inflammatory factors (chemokines, interleukins, TNFα, and complement factors) but their role in brain aging has been studied less^60^. Except for specific genes such as *IL18* in Mic1 from the PFC and *C1QA* and *C1QB* in Mic1 from the hippocampal formation, we did not observe a substantial induction of pro-inflammatory factors (**Figure 5B and S6C**). C1q increases in the hippocampus of old mice^61^, causing synapse removal and delayed OPC differentiation. Interestingly, we noticed a reduction of the TGFβ signaling ligand *TGFB1* in Mic1, especially in the midbrain, hypothalamus, and pons and medulla, although a partial reversion was observed in exceptionally old individuals. TGFβ receptor, *TGFBR1*, was also downregulated in the same regions (**Figure 5F**). TGFβ signaling sustains microglia homeostasis in a cell autonomous manner under physiological conditions, preventing excessive activation of proinflammatory responses^62^. This may explain the pro-inflammatory features of microglia from different regions including the pons and medulla with aging. In this regard, myelin debris reduce *TGFB1* expression^63^. In contrast, the coreceptor *TGFBR2* was upregulated in Mic1 from the hippocampus, striatum, thalamus, and pons and medulla.

Brain ECM not only provides structural support but also stores and activates secreted factors. Accordingly, changes in the composition and organization of the ECM are critically involved in brain aging and neurodegeneration^64^. Collagen is an indispensable component of the brain ECM. Notably, multiple collagen-coding genes such as *COL4A3*, *COL4A4*, *COL9A1*, *COL11A1*, *COL14A1* and *COL21A1* were downregulated in neurons and glial cells, suggesting a rather general disruption of ECM homeostasis during monkey brain aging (**Figure S6C**). We also observed increased tenascin-C (*TNC*) in astrocytes from various non-telencephalic regions. Tenascin-C deposition inhibits myelination in AD by acting on OPCs^65^. Tenascin-R (*TNR*) inhibits myelination in vitro^66^ and was increased in OPCs from different locations.

For signaling factors involved in regulation of myelination (**Figure 5G and S6C**), we identified a broad decrease of contactins (*CNTN1*/*2*) in non-neuronal cells that was more marked in astrocytes and ODCs from the pons and medulla. Contactins are a type of cell adhesion molecules essential for myelin formation and organization^67^. We also noticed that platelet-derived growth factor receptor alpha (*PDGFRA)* and the Wnt signaling coreceptor *LGR5*^14,68^ decreased exclusively in Ast2 and OPCs from the pons and medulla. OPC proliferation and differentiation require PDGF^69^, whereas Wnt/β-catenin signaling has a complex context-dependent role in myelination^70^. SmFISH confirmed the decrease of *PDGFRA* in OPCs from the pons and medulla (**Figure 5H**).

These results emphasize that the molecular changes during cynomolgus monkey brain aging involve complex alterations in multicellular crosstalk, with both general and region-specific patterns.

### Characterization of aging-related cis-regulatory elements

Intricate epigenomic changes underlie aging-related transcriptional alterations in multiple organisms^71,72^. We leveraged the multimodal nature of our dataset to gain insight into the dynamics of these changes in the aging cynomolgus monkey brain.

First, we explored whether epigenetic erosion, defined as a genome-wide reduction of chromatin openness at the TSS despite maintaining a consistent number of total captured open chromatin fragments^73,74^, could explain the observed transcriptional changes in aging. This phenomenon has been used to postulate that drifts in cell identity caused by global epigenetic noise drive transcriptional variation in patients with late-stage AD^73,75^. We observed epigenetic erosion associated with aging in splatter neurons from the hypothalamus and a few non-neuronal cell subtypes (OPCs, ODCs and Mic1) from different regions but with overrepresentation of the hypothalamus (**Figure S7A, B**). The same phenomenon was identified when the analysis was restricted to cell-type specific pDEGs such as myelination-related pDEGs in the pons and medulla (**Figure S7C**). Therefore, epigenetic erosion may contribute to transcriptional changes in selected cell subtypes during monkey brain aging, but there is no evidence for a universal role across all cell subtypes.

Second, we investigated the changes of chromatin accessibility at specific loci with aging. To this aim, we determined the cCREs in each individual cell subtype across all eight brain regions. We identified the open chromatin regions (peaks), which were then combined to generate a union set of 1,907,418 peaks from all cell subtypes. To further refine this list, we calculated the accessibility scores (counts per million, CPM) of these peaks and filtered out those below a threshold (CPM > 4 in at least two individuals in any cell subtype). This yielded a total of 396,892 cCREs enriched in 59 cell subtypes with substantial specificity (**Figure S7D**; *see website*). Of those, 369,110 (93.0%) had orthologous sequences in the human genome (**Figure S7E**). A large proportion of these orthologue sequences (82.5%) overlapped with human brain cell-type specific cCREs defined in a recent multi-brain region snATAC-seq study^72^ and a bulk ATAC-seq dataset from the fetal hippocampus^76^ (**Figure S7F**). The background sequence exhibited a lower proportion of orthologous sequences (75.5%) and, more importantly, a smaller overlap with human brain cCREs (18.9%). Next, we linked the cell-type specific cCREs to putative target genes by integrating them with the snRNA-seq. This yielded a total of 1,346,131 gene-cCRE pairs, 299,269 of which were positively correlated (FDR < 0.05, *R* > 0.3, and < 500 kb to TSS) (**Figure S7G**; *see website*). The latter comprised 111,031 cCREs and 12,306 genes (**Figure S7H**). The genomic distribution of the gene-linked and non-gene linked cCREs included a majority of distal and intronic sequences, but as expected, promoters were more abundant in the former (**Figure S7I**).

We used the refined list of 396,892 cCREs to search for differentially accessible regulatory elements (DAREs) in aging. As with the snRNA-seq, we applied both an age-based linear model (pDAREs) and a pairwise comparison between adjacent age groups (sDAREs) for each cell subtype. This identified 33,954 pDAREs and 51,199 sDAREs, with no obvious bias in terms of nucleus or captured chromatin fragment numbers among the different cell subtypes (**Figure 6A and S7J**; *see website*). There were more sDAREs at the early and late stages than at the very late stage (**Figure S7K**). Moreover, pDAREs and sDAREs with reduced chromatin accessibility upon aging were more abundant than those with increased accessibility (**Figure S7L**). Likewise, as with the DEGs, most DAREs were not shared among different cell subtypes (**Figure S7M, N**). We then evaluated the association of DAREs and their predicted target DEGs, observing a relatively small fraction of DAREs (pDAREs: 10.49%; sDAREs: early stage 9.13%, late stage 6.41%, very late stage 4.63%) associated with changes in gene expression (**Figure S7O**). An interesting representative of the DEG-linked DAREs was the *CAMK2N1* locus, which contained four DAREs exhibiting reduced chromatin openness with aging in upper and deep layer Ex neurons from the PFC (**Figure 6B**). However, the locations of these DAREs differed: in L2/3-IT Ex neurons one DARE was found in the promoter and another in an upstream enhancer, while in L6-CT Ex neurons two DAREs were found in distal downstream enhancer regions. Notably, the cCREs corresponding to these four DAREs were conserved in humans. Other examples of DAREs associated with DEGs included *KIT* in midbrain-derived Inh neurons from the thalamus and midbrain, and *GFAP* in PFC astrocytes (*see website*).

**Figure 6.**
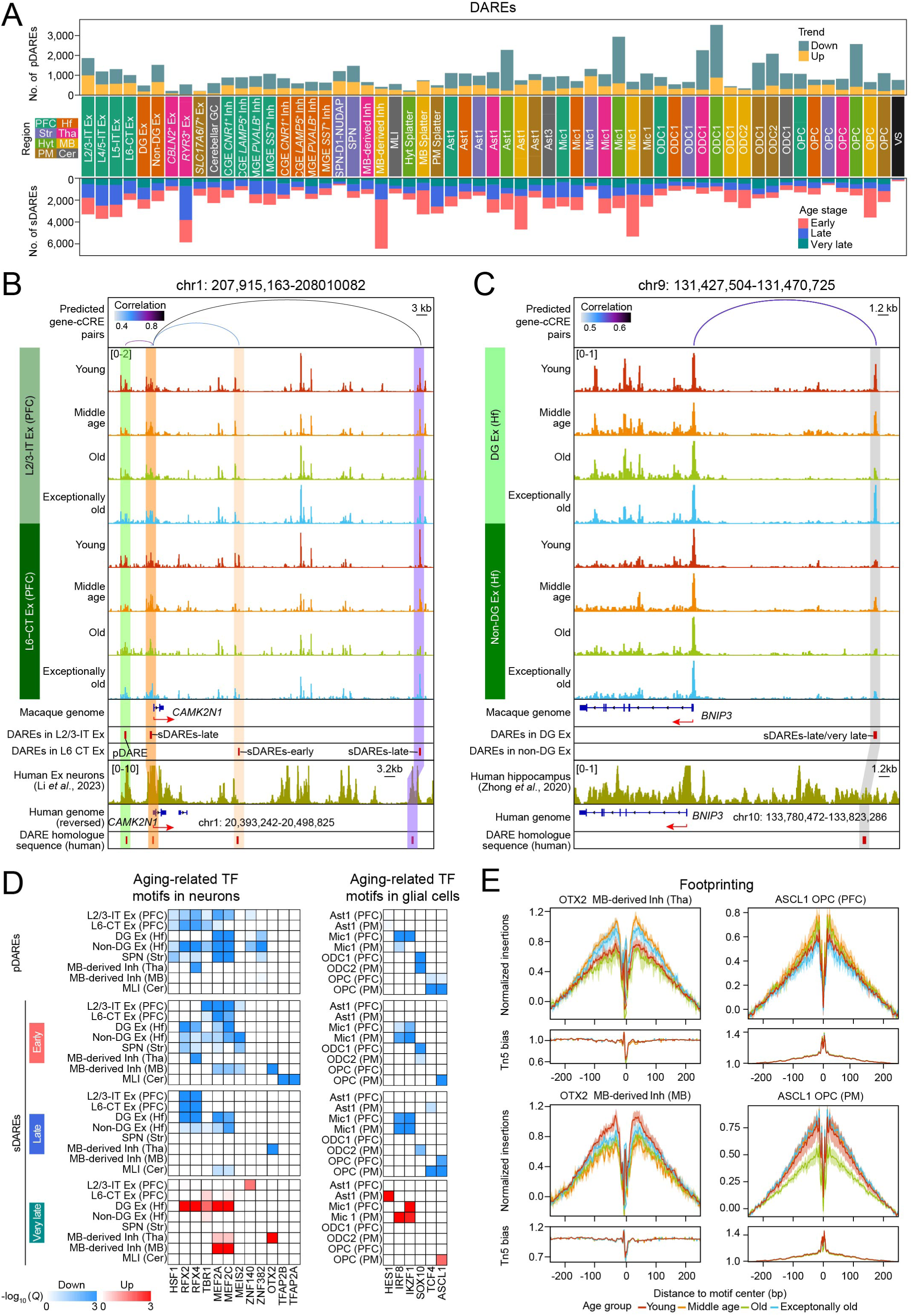
Identification of cell-type-specific *cis*-regulatory elements in brain aging. (A) As in Figure 3A, for pDAREs (top panel) and sDAREs (bottom panel) in 59 cell subtypes. (B) Genome browser tracks showing the normalized aggregate peak signals associated to TSS at the *CAMK2N1* locus for L2/3-IT Ex and L6-CT Ex in the monkey PFC. The degree of association between peaks and genes is represented by the colour scale. Homologous site in humans is displayed in the lower part of the track plot. (C) As in Figure 6B, for the longDARE in *BNIP3* locus from hippocampal DG Ex and non-DG Ex neurons. (D) Heatmap showing the DARE-enriched TF motifs in representative neuronal (left panel) and non-neuronal (right panel) cells. The colour scale indicates the significance of the enriched motif for with aging (downregulated, blue; upregulated, red). (E) Tn5 bias-adjusted TF footprints for OTX2 in midbrain-derived Inh neurons from the thalamus (top left panel) and midbrain (bottom left panel), and ASCL1 in OPCs from the PFC (top right panel) and pons and medulla (bottom right panel), respectively. In each graph, the Tn5 bias-subtracted normalized insertions (y-axis) are plotted from the motif center to 200 bp on either side (x-axis). Abbreviations: DAREs, differentially accessible regulatory elements; pDAREs, progressive differentially accessible *cis*-regulatory elements; sDAREs, stage-specific linear model differentially accessible *cis*-regulatory elements.

We also studied open chromatin regions enriched in exceptionally old monkeys, which we termed longevity-associated DAREs (longDAREs). Intriguingly, DG Ex neurons contained the highest number of these (62), with more than half of the corresponding cCREs conserved in the human brain (**Figure S7P**; *see website*). Among the latter, we noticed a longDARE linked to *BNIP3* (BCL2-interacting protein 3) and another to *SOX9*, both in distal enhancers (**Figure 6C and S7Q**). BNIP3 is a potent inducer of apoptosis and autophagy that was recently associated to brain-regulated longevity in *Drosophila*^77^, whereas SOX9 is a master regulator of neurogenesis^78^.

Our analyses highlight a high degree of cell-type specific complexity in epigenetic and transcriptional regulation in the aging cynomolgus monkey brain, identifying specific loci that may be involved in either brain aging or longevity.

### Aging-related transcription factors

To identify TFs that change with cynomolgus monkey brain aging, we first generated a brain-wide TF profile by analyzing the binding motifs contained in the 396,892 cell-type specific cCREs. After filtering based on expression levels, we identified an enrichment of 198 cell-type specific TFs (**Figure S7R, S**; *see website*). The resulting TFs comprised common and cell-type specific TFs. Common TFs present in most cell subtypes included for example CTCF, which maintains higher-order chromatin structures^79^, the mitochondrial respiration regulator NRF1^80^, and RFX2/4, which are related to ciliogenesis (a function related to neuronal maturation)^81^. Cell-type specific TFs included TBR1 in PFC neurons, MEIS2 in SPNs, OTX2 in midbrain-derived Inh neurons, HES1 in astrocytes, SOX10 in ODCs, ASCL1 in OPCs, and FOXP1/2 in vascular and stromal cells^72^, among others.

Next, to predict the candidate TFs shaping chromatin accessibility in the aging monkey brain, we calculated the enrichment of TF motifs within the identified DAREs (pDAREs and sDAREs). This yielded 199 TFs enriched in the DAREs with reduced chromatin accessibility and 180 in those with increased accessibility. In sDAREs, the number of downregulated TF motifs exceeded that of upregulated motifs at all stages, except for the very late stage. Among the TF motifs enriched in neurons, OTX2 and MEF2A/C motifs were reduced at the late stage in midbrain-derived Inh neurons from the thalamus and Ex neurons from the hippocampal formation, respectively (**Figure 6D**). OTX2 controls the development of different neuronal populations in the midbrain and thalamus and is also expressed in the adult^82^; MEF2 factors are master neuronal regulators of higher-order brain functions such as cognition, memory, and learning^83^. Likewise, motifs for HSF1, a key mediator of stress responses^84^, and RFX2/4, which are involved in neuronal differentiation during development^81^, were broadly reduced in telencephalic neurons. Impaired expression of RFX family TFs has been linked to neuropsychiatric diseases. Among the TF motifs downregulated in glial cells, we noticed a reduction of IKZF1 and IRF8 motifs in the microglia from the PFC and pons and medulla starting at the early stage. IKZF1 regulates microglia homeostasis^85^, whereas IRF8 is a critical regulator of microglia-specific chromatin accessibility and DNA methylation^86^. In addition,

ASCL1 and TCF4 motifs were downregulated in OPCs from the pons and medulla, albeit with different kinetics, the former at the early stage and the latter at the late stage. ASCL1 is a well-known regulator of OPC differentiation into ODCs^87^ and TCF4 a major effector of Wnt signaling^88^. These findings further support that the pons and medulla is a major hotspot for multicellular network dysfunction contributing to myelination injury in the aging monkey brain. The enrichment for TFs such as OTX2 in midbrain-derived Inh neurons from the thalamus, RFX2/4 and MEF2 factors in DG Ex neurons from the hippocampal formation, and ASCL1 in OPCs from the pons and medulla was reversed at the very late stage. The loss and reacquisition of binding motifs for these TFs were confirmed by footprinting analysis (**Figure 6E**). Moreover, we noticed that the Notch downstream effector HES1, whose activity did not change at other life stages, was selectively increased at the very late stage in Ast1 from the pons and medulla. HES1 promotes the choice of astrocyte versus OPC differentiation in gliogenesis during development^89^, a phenomenon that is maintained at low rate in the adult brain^90^. This suggests that reactivation of developmental programs helps overcome some of the detrimental effects of monkey brain aging.

This dynamic repertoire of TFs and their targets should help understand the crosstalk between transcriptional and epigenetic changes in the aging cynomolgus monkey brain.

### Association between risk variants of human diseases and monkey brain aging

We envisaged that our dataset could be useful to study the interplay between primate brain aging and human heritable risk variants associated with high-order brain functions or neurological diseases.

To this end, we performed linkage disequilibrium score regression (LDSC) analysis^91^ between the 369,110 cynomolgus monkey cell-type specific cCREs whose sequence is conserved in human and a panel of GWAS variants (**Figure 7A and Table S4**). We included cCREs irrespective of whether the chromatin accessibility peak had a match in the combined human brain snATAC-seq dataset. As expected, ‘Intelligence*’* and ‘Memory performance’ were broadly associated with neurons, and ‘Multiple sclerosis’ and ‘Alzheimer’s disease’ with microglia. Stronger associations were observed at conserved peaks. Two traits related to cognition, ‘Cognitive G4’ and to a much lesser extent ‘Cognitive G6’^92^, were exclusively enriched in conserved cCREs but showed different cell-type specificity. ‘Cognitive G6’, related to reaction time and attention, was enriched in L2/3 Ex neurons from the PFC, whereas ‘Cognitive G4’, representing intelligence and short-term memory, was enriched across multiple brain regions but particularly in molecular layer interneurons from the cerebellum and *CBLN2*^+^ Ex and *RYR*^+^ Ex neurons from the thalamus. This supports specialized roles of various neuronal populations in cognitive functions. Interestingly, ‘White matter integrity’ was associated with OPCs and ODCs at non-conserved peaks, and with astrocytes and ODCs at conserved peaks.

**Figure 7.**
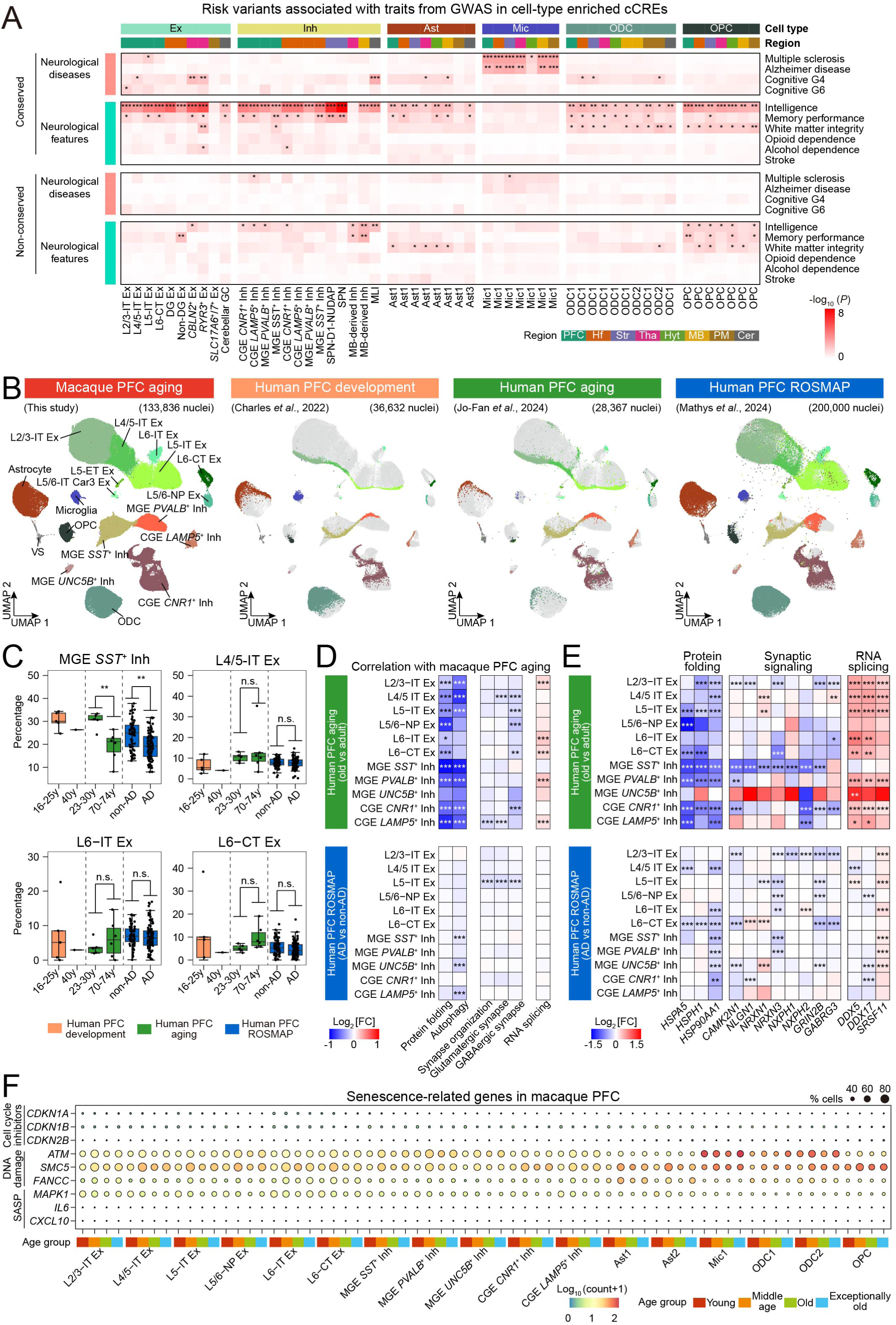
Comparison of monkey aging with human aging and Alzheimer’s disease. (A) Heatmap showing the enrichment of risk variants associated with neurological diseases and neurological features from GWAS studies in the conserved (top panel) and non-conserved (bottom panel) cell-type specific cCREs. Cell subtype and region identity are shown on the top. (B) Mapping of single-cell data of the PFC from human development^1^, aging^12^ and AD^4^ datasets to the monkey PFC by label transferring. (C) Box plots comparing the proportion of the indicated cell subtypes in the PFC from human development, aging, and AD. The dots indicated each individual. (D) Heatmap showing the fold change (log_2_) value for different gene signatures (grouped as score, *see website*) related to monkey brain aging in neuronal subtypes comparing old and young adult PFC (top panel), and AD versus non-AD PFC (bottom panel). (E) As in panel (**D**), for the fold change (log_2_) value for a panel of genes related to protein folding, synaptic signaling, and RNA splicing. (F) Bubble plot showing the expression levels for representative cellular senescence-associated genes in the categories of cell cycle inhibitors, DNA damage, and SASP for each cell subtype within the monkey PFC.

To gain further insights, we applied LDSC analysis between GWAS variants and DAREs (pDAREs and sDAREs) (**Figure S8A**; *see website*). There was association between DAREs in diverse cell subtypes and brain regions with ‘Intelligence’, without a noticeable pattern. DAREs in astrocytes from the midbrain and pons and medulla, as well as in cerebellar granule cells, CGE-derived *CNR1*^+^ Inh neurons from the PFC, and MGE-derived *LAMP5*^+^ Inh neurons from the hippocampal formation, were associated with AD. However, AD traits were overall more substantially associated with DAREs in microglia from different monkey brain regions. Likewise, MS traits were associated with DAREs in neurons from telencephalic regions, midbrain and pons and medulla, and microglia from the midbrain and hypothalamus^93^. We then focused on GWAS lead variants overlapping with DAREs. Interestingly, a DARE with increased chromatin accessibility with aging in the microglia from the pons and medulla overlapped with AD risk variant *rs1870138* in *TSPAN14* (**Figure S8B**). This variant impairs *TSPAN14* gene expression by reducing the binding of the TF TAL1 at a putative enhancer^94^. Likewise, rs76514293 and rs77226570 in *TMEM87B*, which has been linked to MS^95^, were associated with microglia from the midbrain (*see website*).

These results support the potential of our dataset for investigating the functional consequences of human genetic risk factors in the context of aging-induced chromatin changes in the cynomolgus monkey brain. They also underscore interspecies differences that should be considered when using NHPs for human disease modeling.

### Shared molecular features of primate brain aging and human disease

Above we have shown that cynomolgus monkey brain aging exhibits specific molecular features shared with human neurodegeneration, particularly AD. To investigate more systematically the differences between primate brain aging and AD, we conducted a comparative analysis between our monkey PFC dataset and publicly available snRNA-seq datasets of the human PFC. The latter included two recent studies from young and old adults (16-74 years old) without signs of neurodegeneration^1,12^, and another study with AD patients compared to age-matched controls from the ROSMAP consortium^4^. By projecting the human datasets onto our snRNA-seq UMAP using orthologous genes for clustering, we observed good consistency in cell subtype annotations across all datasets (**Figure 7B**). Furthermore, analysis of cell proportions revealed a consistent reduction in MGE *SST^+^* Inh neurons in older humans, regardless of AD status (**Figure 7C**). However, no significant changes were observed in the proportions of L6-IT Ex and L6-CT Ex neurons.

We next investigated whether pDEGs in monkey PFC cell subtypes exhibit similar trends in human aging and AD. For neurons, multiple downregulated pDEGs related to protein folding and synapse regulation, as well as upregulated genes associated with RNA splicing, showed similar patterns in both monkey and human brain aging (**Figure 7D, E**). A similar trend was observed for the former two functional categories in AD, albeit the changes were less pronounced. In contrast, RNA splicing-related genes displayed a substantially different pattern in AD, with many either unchanged or exhibiting the opposite trend compared to monkey neurons. We also investigated glial cells. Because the number of glial cells captured in the aging human brain was not sufficiently balanced across donors^12^, we focused on AD PFC and their age-matched controls^4^. No obvious concordance was observed between both species except for HIF1 signaling in ODCs (**Figure S8C**).

It has been proposed that senescent cells contribute to the pathogenesis of AD^96,97^. Senescence is characterized by the upregulation of cell cycle regulators, DNA damage response factors, and tissue-damaging proinflammatory factors. We observed no clear overall enrichment of a panel of representative genes associated with senescence in neurons or glial cells from the PFC of either old monkeys or humans, including AD patients (**Figure 7F and S8D, E**). However, DNA damage response-related genes (*ATM*, *SMC5*, *FANCC*)^98^, and *MAPK1* (linked to the senescence-associated secretory phenotype)^99^ increased in primate aging and AD, more apparently in glial cells. Our findings highlight the value of NHPs as models for studying human brain aging. They also underline that it is an oversimplification to regard neurodegeneration in AD as an exacerbation of normal aging processes, as the mechanisms at play are distinct.

## DISCUSSION

Our dataset represents the largest cell atlas of NHP brain aging to date and provides a foundation for further expansion, incorporating additional cynomolgus monkey brain regions and conditions. Although selected NHP brain regions at different ages had been profiled with sn-RNA-seq in previous studies^10,11,15,16^, these studies do not involve multiple regions from the same individuals, lack chromatin accessibility data, and do not include exceptionally old individuals. Moreover, biases in cell proportions due to sampling in those datasets limit the ability to analyze differences in specific cell populations across the lifespan.

The transcriptional analyses highlight a series of novel points with potentially relevant implications (**Figure S8F**). For example, we demonstrate that *SST*^+^ Inh neurons^4,13^ and L6 Ex neurons in the PFC have an enhanced susceptibility to death across monkey lifespan compared to other neurons in this region. *SST*^+^ Inh and L6 Ex neurons play relevant roles in sleep homeostasis, mood control and sociability^100,101^. Notably, in contrast to *SST*^+^ Inh neurons, L6 Ex neurons decline in middle-aged monkeys. This raises the question of whether this phenomenon reflects prolonged brain maturation in adult primates rather than degeneration associated with aging^1^. Meta-analysis of previously reported human datasets did not show a reduction in L6 Ex neurons with aging and similar conclusions were made in a recent scRNA-seq study of human brain maturation^1,12^. Nevertheless, it should be noted that those studies include few middle-aged individuals and suffer from high interindividual variability. Regarding potential underlying mechanisms, an increase in a classical senescence signature with aging was not observed at the transcriptional level in any cell subtype from the PFC. We observed, however, increase in DNA damage response genes with aging, particularly in glial cells. We cannot exclude that classical senescence is primarily manifested at the protein level or that the newly appearing senescent cells are scarce and thus were missed in the single-cell analysis^102^. Experiments such as single-cell ribosome profiling^103^ and high-resolution spatially resolved transcriptomics^19^ will help elucidate these possibilities. We also show that cynomolgus monkey brain aging affects fundamental processes such as synapse regulation and myelination. These functions have long been recognized to be altered with aging in mammals including primates^104,105^, but the causes are not well understood. We reveal that a variable mixture of general, region-specific, and cell-specific mechanisms underlies these changes. Region-specific mechanisms affecting neurons are more frequent in the telencephalon, particularly in the PFC, which may reflect specific microenvironmental features. In this regard, we provide evidence that PFC Ast2 are more vulnerable to aging than other astrocyte subtypes in any of the regions tested. Furthermore, we identified the pons and medulla, located at the intersection of axonal bundles connecting the upper brain, cerebellum, and spinal cord, as a major target area for aging-induced multicellular dysfunction leading to myelination injury. This is consistent with existing knowledge that mouse brain regions rich in or surrounded by white matter are more severely affected by aging^7^. We also uncovered altered pathways involving growth factors, inflammatory responses, and extracellular matrix organization in other cell subtypes and regions. Collectively, these findings suggest that therapies aimed at promoting healthy brain aging in primates should target multiple cell types and pathways simultaneously to be effective.

The chromatin accessibility analysis also yielded interesting observations (**Figure S8F**). Importantly, only a proportion of aging-related differentially accessible peaks in the snATAC-seq dataset, defined as DAREs, correlated with changes in gene expression. This may be explained by the fact that long-range chromatin interactions, often involving transcriptional factories at the junction of distant chromosomal regions, play a role in regulating transcription^106^. Such scenario reinforces the idea that progressive and pervasive chromatin dysfunction may be a driving engine of primate brain aging^75^. The observation of epigenetic erosion in selected monkey brain cell subtypes with aging supports this idea. Performing assays such as high-throughput chromosome conformation capture^107^ will help clarify this. Other layers of gene regulation such as DNA methylation, histone modifications, transcriptional pausing or mRNA stability may also contribute to the discrepancies between gene expression and chromatin accessibility^107^. Regardless, we have identified potentially relevant associations such as the *CAMK2N1* locus, where DAREs in different locations likely determine changes in gene expression with aging in PFC Ex neurons.

In addition, the characterization of the repertoire of TFs involved in monkey brain aging provided noteworthy insights. This included supporting evidence for the increased vulnerability of the multicellular networks from the pons and medulla and the reversal at the very late stage for multiple TFs reduced at earlier stages. The latter correlates with the observation that many DEGs are partially reversed in multiple cell subtypes and brain regions in exceptionally old monkeys that possibly represent elite survivors. Whether the resilience to perturbation in those monkeys is a result of brain crosstalk with body functions and how this may happen will require further investigation. Notably, there were also specific changes in chromatin accessibility, defined as longDAREs, in exceptionally old monkeys, which, intriguingly, were more enriched in DG Ex neurons. The extent to which neurogenesis in the hippocampal DG contributes to neuronal turnover in adult primates is under debate^21^. Nevertheless, mature neurons in this area play fundamental roles in new memory formation, which is severely impaired with aging. Interestingly, among the genes corresponding to these longDAREs, we observed *BNIP3*, which has been linked to longevity in flies through unclear mechanisms^77^. Besides, we established associations between monkey DAREs in different cell subtypes, more obviously microglia, and human risk variants for AD or MS in GWAS. An example of DAREs associated with AD risk variants in GWAS was the *TSPAN14* locus. This risk variant reduces the expression of the surface protease ADAM10 and has been connected to human microglia activation in AD^94^. Likewise, we performed a systematic comparison of our PFC snRNA-seq dataset with previously reported human datasets from the PFC of young versus aged individuals and AD patients^1,4,12^. Aging-related processes exhibited considerable transcriptional similarities between both types of primates. However, while there were commonalities, we also observed substantial differences between brain aging and AD.

Collectively, we have generated a vast dataset of cynomolgus monkey brain aging difficult to generate with other primates like human. Our work can be explored interactively through our dedicated online portal (https://db.cngb.org/stomics/nhpabc/), which is designed to facilitate the generation of new observations and hypotheses. We have provided comprehensive analytical pipelines; however, the use of different algorithms, including machine learning approaches, could yield new insights. This atlas, along with its future expansions, may serve as a valuable resource for developing targeted interventions to promote healthy brain aging and longevity in primates. Such treatments could be initially tested in NHPs, with outcomes assessed both clinically and experimentally using the multimodal omics approach presented here.

### Limitations of our study

Increasing the number of nuclei will further improve statistical power and enable more detailed assessments of cell trajectories in aging. Similarly, including additional brain regions will offer a more comprehensive understanding of monkey brain aging. To ensure homogeneity we selected female monkeys, but sex influences longevity and the susceptibility to neurodegeneration in older primates^108,109^. Thus, the observed phenomena or their kinetics may differ in male monkeys. Likewise, there may be differences between NHP species and those that are evolutionarily closer may be more similar. It is also important to note that we cannot determine whether the monkeys sacrificed at an earlier stage could have reached an exceptionally old age. However, exceptional longevity in farmed colonies is rare^18^, implying that the probability of a younger monkey achieving the oldest ages studied here is low. This supports the possibility that the molecular events associated with aging may have had a different trajectory in exceptionally old monkeys. Regarding the changes in cell proportions, it should be considered that cell capture using droplet-based snRNA-seq approaches does not necessarily reflect the true cell composition of the tested tissue. Future work involving a larger cohort of monkey samples across all life stages, along with the use of additional cell quantification methods, will help clarify this. As for the comparisons between monkey brain aging and human neurodegeneration, cynomolgus monkeys and humans may not exhibit the same molecular features nor the underlying processes have the same dynamics.

## METHODS

### Ethical statement

Cynomolgus macaques in this study were group-housed at Guangdong Landau Biotechnology Co., Ltd (Guangzhou, China). The facility has been fully accredited by the Association for Assessment and Accreditation of Laboratory Animal Care International (AAALAC) since 2011. All experimental procedures were reviewed and approved by the Institutional Animal Care and Use Committee of Guangdong Landau Biotechnology Co., Ltd (LDACU-20201216-01) and the Institute of Review Board of Bioethics and Biosafety of BGI (BGI-IRB A21022). Studies were implemented in compliance with the National Institutes of Health (USA) Guide for the Care and Use of Laboratory Animals (8th edition, 2011), as well as the ARRIVE guidelines (https://arriveguidelines.org/) for reporting animal research. Monkeys were fed with a commercial diet (Ke-Ao, HFZ-15kg) twice daily, along with disinfected seasonal fruits and vegetables once daily. Fresh water was available *ad libitum*. Daily monitoring of behavioral states was conducted by the husbandry staff and veterinarians.

### Dissection and collection of monkey brain tissue

Thirteen healthy female cynomolgus monkeys ranging in age from 5 to 29 years old were selected for this study. Monkeys were first anesthetized with ketamine (2 mL, 0.1 g/2 mL, intramuscular injection; Zhongmu) and then pentobarbital sodium (20 mg/kg, intramuscular injection; Sigma, P3761) for euthanasia. Brains were perfused with sterile saline solution and extracted from the cranium after euthanasia. PFC, hippocampal formation, striatum, thalamus, hypothalamus, midbrain, cerebellum, and pons and medulla were carefully isolated and placed on an ice-cold board for dissection. Each tissue was cut into 5–10 pieces of roughly 50–200 mg each. All procedures were done within 1 hour. Samples were transferred to cryogenic vials, flash frozen in liquid nitrogen and stored in liquid nitrogen until nuclear extraction was performed.

### Tissue processing and nucleus isolation

Nucleus isolation was conducted following a previously established protocol^110^. Briefly, the tissue was homogenized in a 2-mL Dounce grinder (Sigma, D8938) using 30 strokes with pestle A in 1.5 mL of homogenization buffer. The homogenization buffer consisted of 250 mM sucrose, 10 mg/mL BSA, 5 mM MgCl_2_, 0.12 U/μL RNasin (Promega, N2115), and 1× cOmplete Protease Inhibitor Cocktail (Roche, 11697498001). Following filtration through a 70-μM cell strainer, the mixture was supplemented with an additional 750 μL of homogenization buffer and further homogenized with 15 strokes using pestle B. The resulting suspension was filtered through a 30-μM strainer and subjected to centrifugation at 500 *g* for 5 minutes at 4 °C. The nuclear pellet was resuspended in 5 mL of nucleus wash buffer, which is the homogenization buffer supplemented with iodixanol (Sigma, D1556) at a final concentration of 20%, and then centrifuged at 800 *g* for 10 minutes at 4 °C. This wash and centrifugation step was repeated twice. Finally, the purified nuclei were resuspended in PBS containing 0.04% BSA and divided into two aliquots for subsequent snRNA-seq and snATAC-seq library preparation.

### snRNA-seq library preparation and sequencing

Nucleus suspensions and barcoded beads were loaded onto the droplet generation chip to generate droplet emulsions. Bead collection, reverse transcription, cDNA amplification, and 3’ library construction were performed according to the manufacturer’s instructions. snRNA-seq libraries for the PFC, hippocampal formation, striatum, thalamus, and cerebellum were prepared using the DNBelab C Series Single-Cell Library Preparation Set (MGI, 1000021082) and the hypothalamus, midbrain and pons and medulla using the DNBelab C Series high-throughput Single-cell RNA Library Preparation Set V2.0 (MGI, 940-000519-00). Library concentrations were measured using Qubit ssDNA Assay Kit (Thermo Fisher Scientific, Q10212). The libraries were sequenced on the DNBSEQ-T1 sequencer at the China National GeneBank (Shenzhen, China) with the following sequencing read lengths: for the snRNA-seq libraries of PFC, hippocampal formation, striatum, thalamus and cerebellum: 41-bp read length for read 1 and 100-bp read length for read 2; for the hypothalamus, midbrain and pons and medulla: cDNA 30-bp read length for read 1 and 100-bp read length for read 2, droplet barcode 20-bp read length for read 1 and 30-bp read length for read 2.

### snATAC-seq library preparation and sequencing

Nuclei were extracted using the same protocol described above. After Tn5 tagmentation, transposed nucleus suspensions were converted to barcoded snATAC-seq libraries through droplet encapsulation, pre-amplification, emulsion breakage, captured bead collection, DNA amplification and purification. Indexed libraries were prepared according to the manufacturer’s protocol. snATAC-seq libraries were prepared using the DNBelab C Series Single-Cell ATAC Library Preparation Set (MGI, 1000021878). Library concentrations were measured with a Qubit ssDNA Assay Kit. Libraries were sequenced on a DNBSEQ-T1 sequencer at the China National GeneBank with the following sequencing strategy: 50-bp read length for read 1 and 76-bp read length for read 2.

### Single-molecule fluorescence in situ hybridization (smFISH)

OCT-embedded sections, with thicknesses of 16 μm for the PFC and 20 μm for pons and medulla, were stained using the RNAscope Fluorescent Multiplex and RNAscope Multiplex Fluorescent V2 kits (Advanced Cell Diagnostics, cat. no. 323100) following the manufacturer’s instructions with slight modifications: the target retrieval step was conducted with a boiling time of 15 minutes, and the incubation with Protease Plus was performed at 40 °C for 30 minutes. Nuclei were counterstained with DAPI at a concentration of 0.25 μg/ml (Life Technologies, S36964). The probes for the target genes are detailed in **Table S5**. Images of the PFC slides were taken using a LSM 800 microscope (ZEISS), while images of the pons and medulla sections were acquired with a Tissue FAXS Plus ST (TG). The number of L6 Ex neurons (*SEMA3E^+^/SLC17A7^+^*) was counted manually with ImageJ (v1.54f). Five randomly chosen fields from the deep layer region of each section were included for calculation, with two sections per individual (two individuals per age).

### snRNA-seq data preprocessing, clustering, and cell-type annotation

Raw sequencing reads were filtered, demultiplexed, and aligned to Macaca_fascicularis_5.0 reference genome using a custom workflow (https://github.com/MGI-tech-bioinformatics/DNBelab_C_Series_HT_scRNA-analysis-software)^14^. Both exonic and intronic reads were counted for alignment because of the large amount of unspliced pre-mRNA in the nucleus. Nuclei with number of detected genes per nucleus >500 and a percentage of mitochondrial genes <1% were kept for further analysis. Doublets were identified and removed using DoubletFinder (v2.0.3) with default settings. The Seurat package (v4.2.0)^111^ within an R environment (v4.1.3) was utilized for snRNA-seq downstream analysis. Briefly, the filtered matrix was merged and normalized using SCTransform-based methods. Dimensionality reduction was achieved through principal component analysis (PCA) of the top 3,000 highly variable genes. Batch effects were corrected using Harmony^112^. UMAP plots were generated with the top 30 principal components with the number of neighbors set at 10. Cell clusters were identified and annotated based on canonical cell markers and reviewed manually. For the reclustering of DG Ex neurons, DA neurons, splatter neurons, astrocytes, oligodendrocytes, microglia, vascular and stromal cells, we applied the same methodology.

### Analysis of cell type composition in snRNA-seq

We calculated cell type proportion changes in individuals across age groups by quantifying the number of cells of a given cell subtype in a specific sample (individual by region) relative to the total number of cells. The method for calculating these proportions varies across different brain regions. Specifically, for the PFC, hippocampus, striatum, and cerebellum, we computed the proportion of Ex neuron subtypes relative to all Ex neurons; the proportion of Inh neuron subtypes relative to all Inh neurons; and the proportion of glial cell subtypes (including astrocyte, microglia, oligodendrocytes and OPCs) relative to all glial cells in each specific sample. In the other four brain regions, we calculated the proportion of neuronal subtypes relative to all neurons and used the same method for glial cells. Once the cell proportions were established, we calculated the Pearson correlation coefficient between cell proportions and individual age. A permutation test was then conducted to assess the reliability of age-related changes in cell type proportions. Cell subtypes with *Q* < 0.05 and an absolute *R* > 0.6 were considered significantly associated with aging.

### Identification of differentially expressed genes (DEGs) in snRNA-seq and functional enrichment analysis

*Identification of progressive aging-related DEGs (pDEGs).* We used a previously published method^24,25^. Specifically, only cell subtypes containing more than 10 nuclei per individual and over 100 nuclei per age group were selected for the analysis, and only protein-coding genes were considered. The dataset was filtered to retain only protein-coding genes. The MAST package (v1.12.0) in R was utilized to perform the analysis, employing a linear model that treated age as a continuous variable. To account for multiple comparisons, the Benjamini-Hochberg correction was applied, with the number of comparisons equal to the number of protein-coding genes. Genes with *Q* < 0.01 and an age coefficient threshold of 0.005 (in the unit of log_2_[fold-change]/year, corresponding to around 10%-fold-change from 5 years to 29 years) were considered as DEGs.

*Identification of stage-specific aging-related DEGs (sDEGs)*. The identification of sDEGs was performed using Seurat. For this analysis, only cell subtypes with more than 100 nuclei in each age group were included, and only protein-coding genes were considered. The comparisons between different age stages were conducted using a hurdle model tailored for scRNA-seq data, implemented in MAST^113^. Genes with *Q* < 0.05 and a log_2_[fold change] > 0.25 were considered as DEGs. The same method was used for the analysis of DEGs in human brain aging and AD datasets. To determine the bias in the number of DEGs by the number of nuclei/genes captured, we calculated the Pearson correlation coefficient between the number of pDEGs or sDEGs in each cell type and the number of nuclei/genes captured in the corresponding cell type.

*Functional enrichment.* The DEGs were analyzed using Metascape^114^, with a *Q* value threshold set at 0.05.

### Pairwise comparison of DEGs and analysis of disease-associated signatures

GeneOverlap (v1.8.0) was applied to compare the DEGs identified in different cell types. A Benjamini–Hochberg procedure was applied to control for multiple comparisons, with a *Q* value threshold set at 0.05. The results were plotted as heatmap using pheatmap (v.1.0.12) in R. To map the expression of disease-associated signatures, the corresponding gene sets were acquired from published studies (*see website*). These signatures were mapped to the corresponding cell types as module scores using Seurat. The age coefficient was calculated as described above for pDEGs.

### Cell-cell interaction analysis

CellChat (v1.1.0)^54^ was used to identify ligand-receptor interaction on snRNA-seq data for each brain region following the standard procedure. Cell types containing more than 10 nuclei per individual animal and over 100 nuclei per age group were selected for the analysis. To enhance the ligand-receptor database, we compiled interaction pairs from NeuronChat, ICELLNETdb, iTALK, SingleCellSignalR, CellTalkDB, Celltalker, and NATMI database^115–122^. We calculated the overall communication probability among cell clusters using the *computeCommunProb* function at each stage, setting the *truncatedMean* parameter to 0. Interaction pairs with a mean communication probability lower than 0.002 and a *P* value ≥ 0.05 were excluded. If either the ligand or receptor was present in the DEG list, the interaction was considered relevant to aging.

### Comparative analyses with human prefrontal cortex development, aging, and neurodegeneration datasets

Human PFC development^1^ and aging^12^ datasets were retrieved from the Gene Expression Omnibus database under the accession numbers GSE168408 and GSE247988, respectively. The AD PFC dataset (ROSMAP)^4^ was taken from the Synapse database (syn52293417, syn44424170), downsampled to 200,000 nuclei. For the ROSMAP dataset, only samples produced using 10x V3 chemistry were considered, and all V2 samples were excluded for the analysis. The data were mapped to our monkey PFC dataset using the *MapQuery* function implemented in Seurat with orthologous genes. The cell proportions and DEGs were determined using the same method mentioned above.

### Stereo-seq data processing

Data were retrieved from Chen *et al.*^17^. The distribution of cell types was determined based on the expression of specific cell-type marker genes, as follows: MGE-derived Inh neurons, *GAD2*/*ADARB2*; CGE-derived Inh neurons, *GAD2*/*LHX6*; LGE-derived Inh neurons, *GAD2*/*FOXP1/PENK*; midbrain-derived Inh neurons, *GAD2*/*OTX2*; Mic1, APBB1IP/CAMD2; Mic2 *APBB1IP*/*DAB2*; ODC1, *MBP*/*OPALIN*; ODC2, *MBP*/*GSN*; Ast1, *SLC1A2*/*GPC5/GFAP^low^*; Ast2, *SLC1A2/GFAP^high^*; Ast3, *SLC1A2*/*SEPT4*. The normalized expression level of GFAP in astrocytes was categorized as follows: levels lower than 1.5 were considered low (*GFAP*^low^), while levels higher than 1.5 were considered high (*GFAP*^high^).

### snATAC-seq preprocessing, clustering, and cell-type annotation

Raw sequencing reads were filtered, demultiplexed and aligned to the Macaca_fascicularis_5.0 reference genome using PISA (https://github.com/shiquan/PISA)^123^, in which the fragment files for each library were generated for downstream analysis in ArchR (v1.0.2). Cells with TSS enrichment score < 4 and fragment numbers < 3,000 were removed. We calculated the doublet score with the *addDoubletScores* function implemented in ArchR and filtered using the *filterDoublets* function with *filterRatio* set at 2. Next, we created 500-bp genomic tiles and determined the accessibility of each cell within these tiles and applied iterative latent semantic indexing (LSI) dimensionality reduction on those by using the *addIterativeLSI* function in ArchR. The clustering algorithm implemented in Seurat was applied to the LSI dimensions with the resolution set at 0.8. Finally, the resulting cell clusters were annotated using the same markers of the snRNA-seq dataset.

### Integration of snRNA-seq and snATAC-seq data

For the integration of global snRNA-seq and snATAC-seq datasets, each was randomly downsampled to 100,000 cells. Log-normalized and scaled gene activity scores served as surrogates for gene expression in cells profiled by snATAC-seq. The *FindIntegrationAnchors* function implemented in Seurat was utilized for data integration, using the intersection of the top 3,000 highly variable genes from each modality as input. Canonical correlation analysis was applied, and the number of neighbors for data integration was set to 10. The same methodology was applied to each brain region but without downsampling.

### Epigenetic erosion

We computed the mean TSS enrichment score for each cell type in each monkey. Then, we calculated the Pearson correlation coefficient between the mean TSS enrichment score and the individual’s age. To assess the reliability of age-related changes in TSS enrichment, a permutation test was conducted. Cell types with *Q* < 0.05 and an absolute *R* > 0.6 were considered significantly associated with aging.

### Identification of candidate *cis*-regulatory elements (cCREs)

*Identification of cell-type enriched cCREs.* Peaks were first identified on the basis of cell subtypes in each individual animal using *addReproduciblePeakSet* function implemented in ArchR, in which MACS2 (v2.2.7.1) was applied^124^. This generated a union peak set consisting of 1,907,418 peaks. To further refine this list, we calculated the CPM of these peaks and filtered out those below a threshold (CPM > 4 in at least two individuals in any cell type). Cell types containing more than 10 nuclei per individual animal and over 100 nuclei per age group were selected for analysis. This yielded a total of 759,236 cCREs enriched in 59 cell types with substantial cell type specificity.

*Generation of peak-to-gene links.* In order to further predict enhancer-promoter interactions in each cell type for each individual brain region, we applied a previously published method^3^ with some adjustments. In brief, co-accessible regions were identified for all cCREs in each cell subtype using *addCoAccessibility* function in ArchR with the following parameters: aggregation k = 10, window size = 500 kb, distance constraint = 250 kb. To determine an optimal co-accessibility threshold for each cluster, a random shuffled cCRE-by-nuclei matrix was used as a background. The distribution of co-accessibility scores from this background was fitted to a normal distribution model. Co-accessibility pairs were tested with a significance threshold of *Q* < 0.01. We then calculated Pearson correlation coefficient between gene expression and cCRE accessibility across matched snRNA-seq and snATAC-seq cell subtype. To do so, we first aggregated all nuclei from snRNA-seq or snATAC-seq for every joint cluster to calculate the average expression levels or accessibility scores. The Pearson correlation coefficient was calculated for every gene-cCRE pair within a 500 kb window centered on the TSS for every gene. We also generated a set of background pairs by randomly selecting regions from different chromosomes and shuffling of cluster labels. Finally, we applied a normal distribution model and defined a cut-off at Pearson correlation coefficient score with an empirically defined significance threshold of *Q* < 0.01 and *R* > 0.3, thereby identifying significantly positively correlated cCRE-gene pairs.

*Identification of conserved cCREs.* Monkey cCREs were converted to the human genome (hg38) using liftOver (v362). Those successfully converted to the human genome were considered to have orthologous sequences in humans. To determine if these cCREs were conserved in the corresponding cell types, we obtained cCREs identified in a recent snATAC-seq dataset of multiple human brain regions^3^, as well as hippocampus bulk ATAC-seq data^76^, and performed a bedtools (v2.31.0) intersection analysis. Monkey cCREs that overlapped with human cCREs were considered as cell-type conserved. Meanwhile, random background sequences were generated from the *Macaca fascicularis* genome, and the same procedures were applied to these sequences for comparison.

### Identification of differentially accessible regulatory elements

*Identification of progressive differentially accessible regulatory elements (pDAREs).* We first aggregated the accessibility scores for each cell-type enriched cCREs across all animals to compute the average accessibility scores for every cell type. To calculate the fold changes in cCREs with aging, we applied the same method used for pDEGs, we then calculated the Pearson correlation coefficient between these average accessibility scores and the age of the animals. To evaluate the reliability of age-related changes in cCRE accessibility, a permutation test was performed. cCREs were deemed significantly associated with aging if they had a *Q* < 0.05, absolute *R* > 0.5 and an age coefficient greater than 0.005.

*Identification of stage-specific differentially accessible regulatory elements (sDAREs).* We used *getMarkerFeatures* function implemented in ArchR with cell-type enriched cCREs as input, comparing between different age stages. cCREs with a *Q* < 0.05 and a log_2_[fold change] > 0.25 were considered as sDAREs.

*Identification of longevity-associated DAREs (longDAREs).* We used *getMarkerFeatures* function in ArchR to compare the nuclei in the exceptionally old group with the rest of the nuclei for each cell type. cCREs with a *Q* < 0.05 and an absolute log_2_[fold change] > 0.25 were considered as longDAREs.

### Transcription factor motif enrichment

Motif enrichment was performed using the JASPAR2020 catalog^125^. TFs expressed in less than 5% of the cells as determined via the matched snRNA-seq cell clusters were excluded. Peaks were annotated with known motifs via *addMotifAnnotations* function in ArchR with default settings. Appropriate GC matched background peaks for each peak were determined using *getBgdPeak* function with default settings. Next, the *computeEnrichment* function was used for determining the enriched motifs in the chosen sets of peaks compared to their respective background.

### Footprinting analysis

TF occupancy was evaluated using footprinting analysis implemented in ArchR. First, putative binding sites for selectively enriched TF motifs were identified using the *addMotifAnnotations* function in ArchR. This step inferred the locations of potential TF-binding sites based on the detected motif enrichment.

Next, footprinting analysis was performed on these putative binding sites using the *getFootprints* function, which considers the known Tn5 insertion bias to more accurately model the TF-DNA interactions and measure TF occupancy at the predicted binding sites. The results were further plotted using the *plotFootprints* function.

### Association of GWAS traits with monkey brain cell subtypes

We performed linkage disequilibrium score regression analysis using LDSC^91^ (v1.0.1), a method for partitioning heritability from GWAS summary statistics. Further analysis was performed according to the standard workflow (https://github.com/bulik/ldsc/wiki) with the cell-type enriched cCREs and summary statistics of traits as input. The cell-type enriched cCREs were converted to human genome (hg19) using liftOver (v362). The summary statistics file for each trait was downloaded from the GWAS catalogue database or published studies as listed in **Table S5.**

### Data availability

All raw data have been deposited to China National Gene Bank (CNGB) Nucleotide Sequence Archive (CNP0004459). All processed data are available at the NHPABC (https://db.cngb.org/stomics/nhpabc/).

## ACKNOWLEDGMENTS

We thank Dr. Dahai Zhu from the Bioland Laboratory (Guangzhou Regenerative Medicine and Health Guangdong Laboratory) for technical help, Dr. Yan Liu from Nanjing Medical University for experimental advice and the CNGB for their support. This work was supported by the National Key R&D Program of China (2022YFC3400400, 2021YFF0702201 and 2023YFD1802400), Shenzhen Basic Research Project for Excellent Young Scholars (RCYX20200714114644191), Shenzhen Key Laboratory of Single-Cell Omics (ZDSYS20190902093613831), Department of Science and Technology of Guangdong Province (2021ZT09Y007), National Natural Science Foundation of China (92368301, 81873736, 82394422, 32230104 and 32373032), Guangdong Basic and Applied Basic Research Foundation (2021A1515012526), Guangdong Provincial Genomics Data Center (2021B1212100001). D.C.R. is supported by the UK Dementia Research Institute through UK DRI Ltd, principally funded by the Medical Research Council. This paper is part of the Mesoscopic Brain Mapping Consortium.

## AUTHOR CONTRIBUTIONS

X.G., Y.Lai and M.A.E. conceived the idea; Chuanyu Liu, Y.Lai, Mingyuan Liu and M.A.E. supervised the work; X.G., Chuanyu Liu, Y.Lai and M.A.E. designed the experiments; Mingyuan Liu and X.Li provided relevant infrastructural support; Xiao Zhang, Y.Lai and M.A.E. wrote the manuscript; X.G. collected all the samples with the help of D.H., M.Pan, L.C., B. Li, W.Y. and P.Y; Xiao Zhang and G.Lai performed the majority of the experiments with the help of Y.Y., H.Zhang, P.F., Xingyuan Liu and P.G.; Xiao Zhang and G.Lai analyzed the data with the help of W.M.; J.A., J.Li., K.Huang, J.Zuo, L.W., S.S., F.Y., Y.X., Y.Hu., Y.Huang., Y.Lin. and Xiaolan Zhang provided technical support; T.Y., J.C. and Q.Zhong constructed the website; S.Li, Y.Lv, B.Q., X.Q., Q.D., Y.H., Chang Liu, L.H., S.Liu, D.Q., Z.Liu., Y.Zhou, L.Q., C.Li., Y.Zhang, G.V., S.G., I.F.D., M.F., R.P.F., J.M., A.P.H., A.K., P.H.M., D.C.R., Xiaolei Liu, H.T., M.Poo, B.W., L.Liu, Y.G., J.X., X.X. gave relevant advice and/or helped edit the manuscript.

## FIGURE LEGENDS

**Figure S1.** Distribution of main cell types in snRNA-seq and snATAC-seq, related to Figure 1. (A) Bubble plot showing the expression levels of canonical marker genes in the main cell types from snRNA-seq. Colour scale represents the average gene expression, while the dot size indicates the percentage of cells expressing a given marker within the population. (B) Normalized genome browser track profiles for the representative marker genes (columns) for the indicated cell types (rows) in the main cell types from snATAC-seq. (C) Stacked bar plot showing the proportions of captured cell types for each brain region across each individual. The bars are coloured by cell type, and grey bars indicate that the specific region could not be analyzed or collected in that individual.

**Figure S2.** Global transcriptomic and epigenomic profiling of individual monkey brain regions, *related to* Figure 2. (A) UMAP visualization of neuronal cell subtypes from the snRNA-seq (top) and snATAC-seq datasets (bottom) in the PFC, hippocampal formation, striatum, thalamus, hypothalamus, midbrain, pons and medulla and cerebellum. The cell type identities and number of profiled nuclei are indicated in the plots. (B) Left: schematic representation of the anatomical regions in a hemi-brain section (section H5) from a five-year-old monkey using Stereo-seq data taken from Chen *et al.*^17^. Right: spatial visualization of Inh neuron subtypes (represented by indicated marker genes) in the same section as the left panel. Scale bar, 5 mm. (C) UMAP visualization of microglia, ODC, OPC, and vascular and stromal cell subtypes from the snRNA-seq (top) and snATAC-seq datasets (bottom). The representative marker genes used to identify each population are indicated in the plot. (D) Left: schematic representation of the anatomical regions in a hemi-brain section (section H7) from a five-year-old monkey using Stereo-seq data taken from Chen *et al.*^17^. Right: spatial visualization of ODC and microglia subtypes (represented by the indicated marker genes) in the same section as left panel. Scale bar, 5 mm.

**Figure S3.** Identification of shared and cell-type specific aging-associated genes in neurons, *related to* Figure 3. (A) Dot plots showing the Pearson’s correlation (*R*) between the number of pDEGs (left panel) or sDEGs (right panel) and the number of captured nuclei or features. Each dot indicates a cell subtype and is coloured according to the brain region. (B) Bar plot showing the numbers of sDEGs with different temporal expression trends. The temporal expression trends were categorized as stage-specific, stage-opposite, and stage-shared groups, with the colors denoting each respective group. For the stage opposite part, the labels at the top of each bar indicate the comparison group (early, late, or very late). (C) Box plot showing the percentage sDEGs in different comparison groups (early, late, and very late) in the overall set of sDEGs. Each point within the box plot represents a neuronal subtype. Dots and boxes are coloured by age stage. (D) Box plot showing the percentage of downregulated and upregulated pDEGs (left panel) and sDEGs (right panel). Each point within the box plot represents a neuronal subtype. The colour represents the expression trend. (E) Dot plots showing the *Q* value (left panel, y-axis), expression levels (right panel, y-axis) and the fold change (age-coefficient, x-axis) of the overall pDEGs in neuronal subtypes. (F) Bar plot showing the functional enrichment of upregulated sDEGs at the very late stage. (G) As in Figure 3C, for three types of sDEGs. (H) Venn diagram showing the overlap between upregulated sDEGs at the very late stage and widely shared downregulated pDEGs. Overlapping DEGs related to the pathways highlighted in panel (**F**) are labelled. (I) Heatmap showing the age-coefficient of genes related to RNA splicing in all neuronal subtypes. (J) Top: line charts showing the module score of genes related to RNA splicing in all the neuronal cells of the indicated brain regions. Bottom: line charts showing the expression levels of *HNRNPA2B1* in all the neuronal cells of the indicated brain regions. The dots represent each individual. (K) Box plot showing the odds ratio values for the four neuronal clusters in pDEGs and sDEGs. Boxes are coloured by cluster. (L) As in Figure 3F, for three types of sDEGs.

**Figure S4.** Identification of shared and cell-type specific aging-associated genes in neurons, *related to* Figure 3. (A) Left: functional enrichment results of cell-type specific upregulated early sDEGs in L6-CT Ex neurons, with a highlight in red on genes associated with chordate embryonic development. Right: line chart showing the expression level of *EBP41L5* in L6-CT Ex neurons, dots and lines in grey represent trends not statistically significant in other cell subtypes. (B) Line charts showing the expression levels of regionally shared DEGs related to synapse structure (*NLGN1*), synapse receptor (*GRIN2B*) and phosphorus metabolism (*CALM3*) in all the PFC neuronal subtypes from each individual. The dots represent each individual. (C) Line charts showing the expression levels of regionally shared DEGs identified in cluster 1 for all hippocampal neurons from each individual. The dots represent each individual. (D) Venn diagram showing the overlap of downregulated (left panel) and upregulated (right panel) regionally shared pDEGs in cluster 1 and widely shared pDEGs. (E) Left: Venn diagram showing the overlap of downregulated (top panel) and upregulated (bottom panel) pDEGs between MB-derived Inh neurons in the thalamus and midbrain. The numbers of overlapping and non-overlapping pDEGs are labelled. Right: line charts showing the expression levels for *SLC6A15* and *KIT*; the dots represent each individual. (F) Specific pDEGs identified in ICj Inh in the stratum, PLI in the cerebellum, and *SLC17A6*/*7*^+^ Ex in the pons and medulla. Specific genes related to the synapse are labelled. The line charts show the expression levels of selected genes, with the dots and lines in grey representing trends that are not statistically significant among other cell subtypes in the corresponding brain regions. The dots represent each individual. (G) Line charts showing the expression levels of genes related to peptidergic neuron identity (*ADCYAP1*, *GHRH*, *CARTPT* and *OXT*) in the hypothalamus.

**Figure S5.** Identification of shared and specific aging-associated genes in non-neuronal cells, *related to* Figure 4. (A) Dot plots showing the Pearson correlation coefficient (*R*) between the number of pDEGs (left panel) or sDEGs (right panel) and the number of captured nuclei or features. Each dot indicates a cell subtype and is coloured by brain region. (B) Box plot showing the percentage of downregulated and upregulated pDEGs (left panel) and sDEGs (right panel) in glial cells. Each dot represents a cell subtype and the colour represents the expression trend. (C) Dot plots showing the *Q* value (left panel, y-axis), expression levels (right panel, y-axis) and the fold change (age-coefficient, x-axis) of the overall pDEGs in glial cell subtypes. (D) Bar plot showing the number of different temporal expression trends among sDEGs and coloured by the characteristics of these trends (stage specific, stage opposite and stage shared). (E) Histogram showing the frequency distribution of downregulated (top panel) or upregulated (bottom panel) DEGs across different cell subtypes and coloured by the expression trend. The plot indicates the proportion of genes that are shared (identified in at least 10 cell subtypes) or cell-type specific (identified in only one cell subtype). (F) Hierarchical heatmap showing the clustering of all glial cell subtypes based on pDEG similarity. Similarity significance (−log_10_ [odds ratio]) is depicted by the colour scale. Region identity is shown on the side of the plot. (G) Bar plot showing the functional enrichment results for downregulated (left) and upregulated (right) pDEGs in vascular and stromal cells. The dotted box on the right-hand side of the plot indicates representative genes corresponding to the functional terms or pathways highlighted. (H) Line charts showing the expression levels of the rRNA processing gene (*RPL5*), transport across blood-brain barrier (*SLC5A5*), and genes associated with AD (*NLGN1*, *BNC2*)^43^, in vascular and stromal cells from each individual. The dots represent each individual. (I) Heatmap showing the age-coefficient value for module scores of different disease-associated microglia (left) and ODC states (right) in monkey microglia and ODC subtypes. The gene lists are taken from previous studies (*see website*). Abbreviations: DAM, disease-associated microglia; MGnD, neurodegenerative phenotype microglia; WAM, white matter-associated microglia; DOLs, disease-associated ODCs.

**Figure S6.** Expression dynamics of ligand and receptors in the aging monkey brain, *related to* Figure 5. (A) Box plot showing the number of aging-associated interactions per region in each DEG category. Dots are coloured by cell type. (B) Stacked bar plot showing the percentage of interactions in different brain regions according to four interaction types, indicated by different colours, across neuronal and glial cells. (C) Heatmap showing the age-coefficient of representative ligand or receptor genes for the indicated pathways. The legend on the top shows the brain region and cell type identity, while the legend on the left indicates the pathways. The colour scale represents the age-coefficient.

**Figure S7.** Characterization of aging-associated *cis*-regulatory elements, related to Figure 6. (A) Volcano plot showing the correlation between the TSS enrichment score of various cell subtypes and chronological age. The cell subtypes that displayed negative association are indicated. *P* values were calculated using a permutation test. (B) Line charts showing the average TSS enrichment score for the specified cell subtypes from each individual. The dots represent each individual. (C) Occupancy plots showing the normalized chromatin accessibility from snATAC-seq data within ± 2 kb of the TSS for ODC2 in the pons and medulla. These plots include all protein-coding genes (top panel) and myelination-related genes identified from the pDEG list (bottom panel). (D) Heatmap showing the 759,236 cell-type enriched cCREs across 59 cell subtypes. The colour scale represents normalized chromatin accessibility. Region and cell subtype identity are shown on the left-hand side legend. (E) Bar plot showing the proportion of sequences converted to the human genome in random background or cell-type enriched cCREs. (F) Venn diagram showing the overlap between the converted random background (left) or the cell-type enriched cCREs (right), and the cCREs identified in the human brain. (G) Pearson correlation coefficient between the chromatin accessibility of cCREs and the expression levels of the associated genes. Those gene-cCRE pairs that are positively correlated (*R* > 0.3 and *Q* <0.05) are highlighted in red. (H) Heatmap showing the normalized gene expression (top panel) and the normalized chromatin accessibility of linked cCREs (bottom panel) in the corresponding cell subtypes. Cell subtype and region identity are shown on the left-hand side legend. (I) Pie chart showing the genomic distribution of cCREs that are not correlated with gene expression (left panel) and those positively correlated with gene expression (right panel). (J) Dot plots showing the Pearson correlation coefficient (*R*) between the number of pDAREs (left panel) or sDAREs (right panel) and the number of captured nuclei or fragments. Dots are coloured by cell subtypes. (K) Box plot showing the proportion of sDAREs in each comparison group. Each point within the box plot represents a cell subtype. The colour indicates the comparison group. (L) Box plot showing the percentage of upregulated or downregulated DAREs in each comparison group. Each point within the box plot represents a cell subtype. The colour indicates the trend of changes. (M) Histogram showing the frequency distribution of downregulated (top panel) or upregulated (bottom panel) pDAREs across different cell subtypes and coloured by the trend of change. The plot indicates the proportion of pDAREs that are shared (identified in at least 10 cell subtypes) and subtype-specific (identified in only one cell subtype), for both the downregulated (top panel) and upregulated (bottom panel) pDAREs. (N) Bar plots showing the number of specific DAREs (left) and shared DAREs (right). The bars are coloured by the trend of change. (O) Box plot showing the percentage of the DAREs linked to DEGs. Each point within the box plot represents a cell subtype. (P) Bar plots showing the number of conserved and non-conserved longDAREs in the indicated cell subtypes. (Q) Genome browser tracks showing the normalized aggregated signal associated to a longDARE at the *SOX9* locus for DG Ex and non-DG Ex neurons in the monkey hippocampal formation. The association between peaks and genes is represented by the colour scale. The human homologue site is displayed in the lower part of the track plot. (R) Heatmap showing the enrichment of 205 TF motifs in cell-type enriched cCREs. Region and cell subtype identity are shown on the right-hand side legend. (S) Box plot showing the number of enriched TF motifs in cell-type enriched cCREs for each cell subtype. Each point within the box plot represents a cell subtype.

**Figure S8.** Additional comparisons of monkey aging with human aging and Alzheimer’s disease, *related to* Figure 7. (A) Bubble plot showing the enrichment of risk variants associated with AD traits from GWAS studies in the DAREs. Cell subtype and region identity are shown on the left-hand side legend. (B) Genome browser tracks showing the normalized aggregated signal at the *TSPAN14* locus in microglia from the monkey PFC (top panel) and pons and medulla (bottom panel). The association between peaks and genes is represented by the colour scale. The DAREs, the corresponding human homologue sites, and SNPs are displayed in the lower parts of the track plot. (C) Heatmap showing the fold change (log_2_) value for different gene signatures (grouped as score, *see website*) related to monkey brain aging in glial subtypes comparing AD and non-AD PFC. (D) Bubble plot showing the expression levels for representative cellular senescence-associated genes in the categories of cell cycle inhibitors, DNA damage, and SASP for each cell subtype within the human PFC comparing young and old adults. (E) As in panel (**D**), comparing the AD and non-AD human PFC. (F) Top left: schematic outlining the study design. Top right: aging selectively reduces MGE-derived *SST*^+^ Inh and L6 Ex neurons in the PFC. Middle: Analysis of dynamic gene expression changes uncovers both common and cell-type specific changes, especially within multicellular networks engaged in synaptic signaling, ECM composition, and TGFβ signaling. An enhanced dysfunction in myelination was observed in the pons and medulla. Bottom left: loci regulated by aging that are connected to longevity and neurodegeneration; changes in these loci are associated with TFs. Bottom right: some aging-associated changes are partially restored at exceptionally old ages and include regulatory loci potentially linked to longevity.

