## Supplementary figures and images for "Multimodal brain cell atlas across the adult macaque lifespan"

### Supplemental Figure

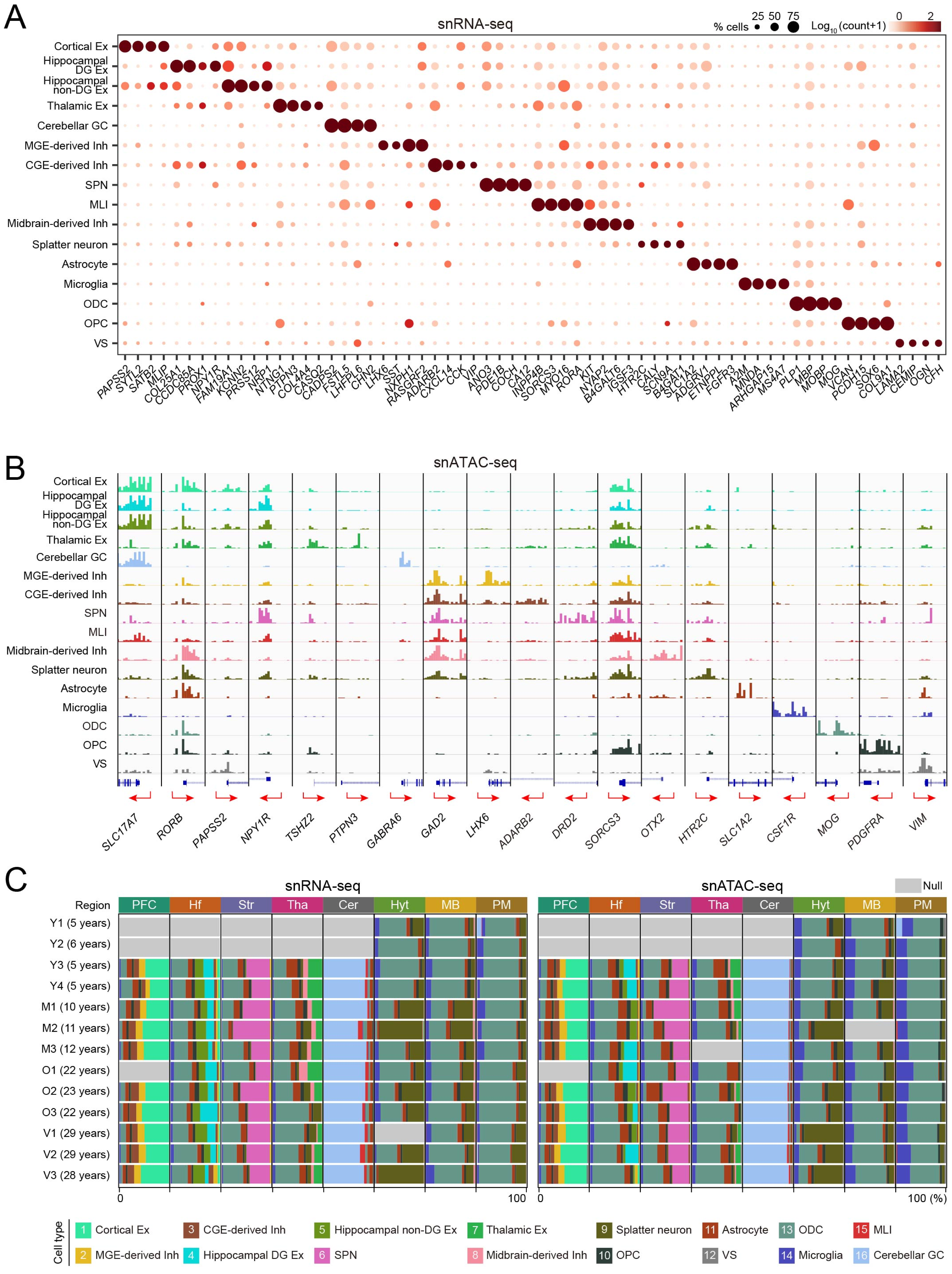

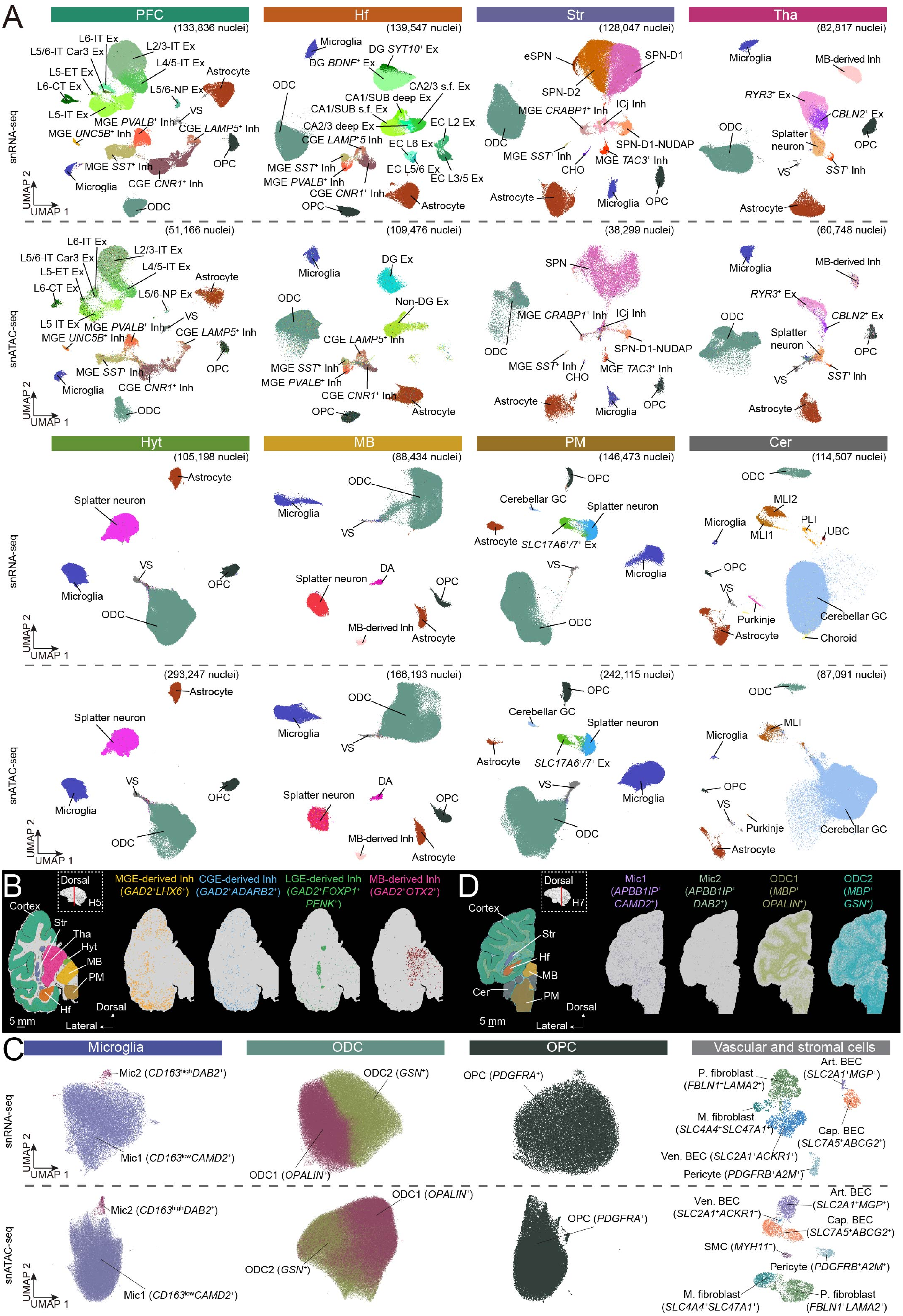

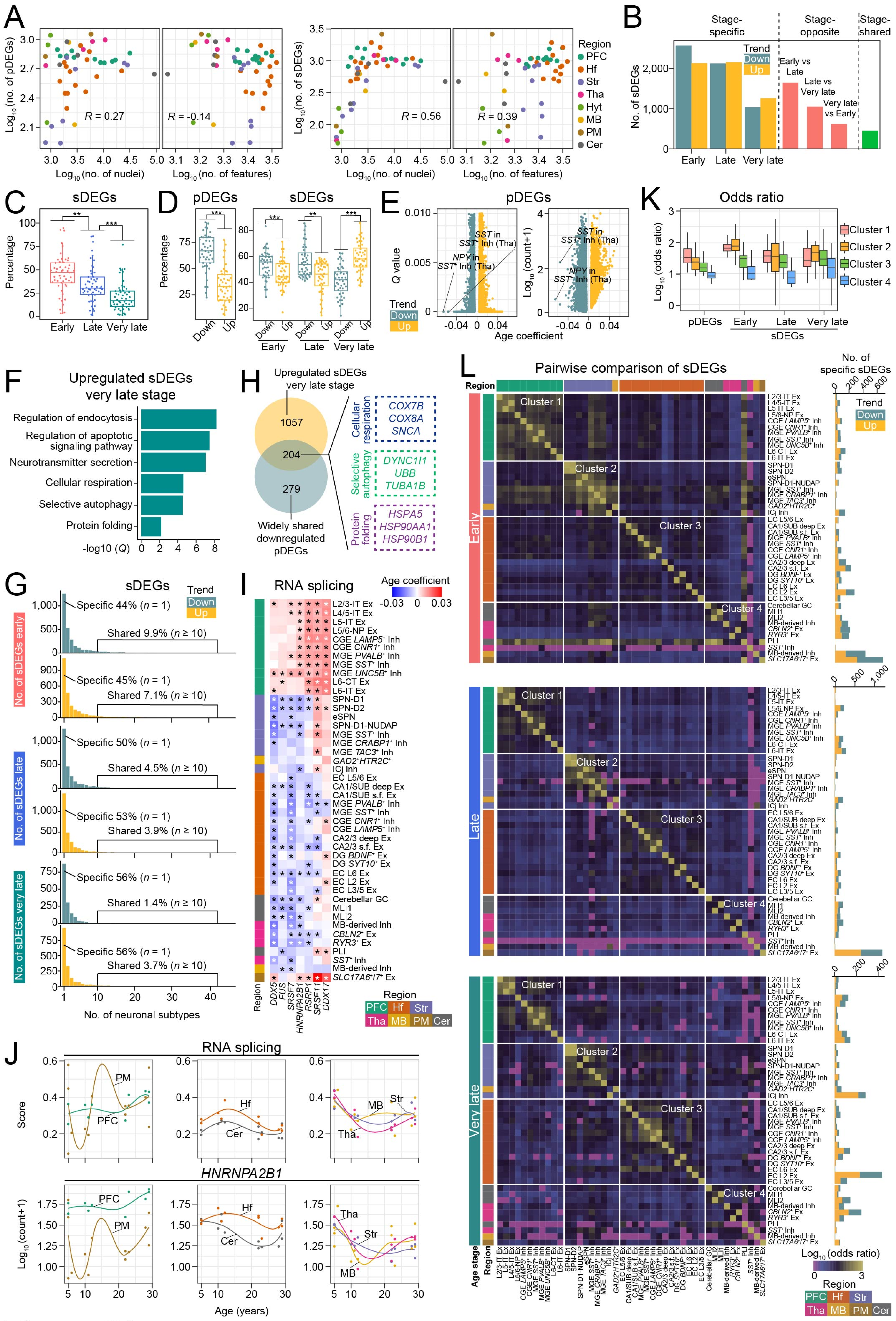

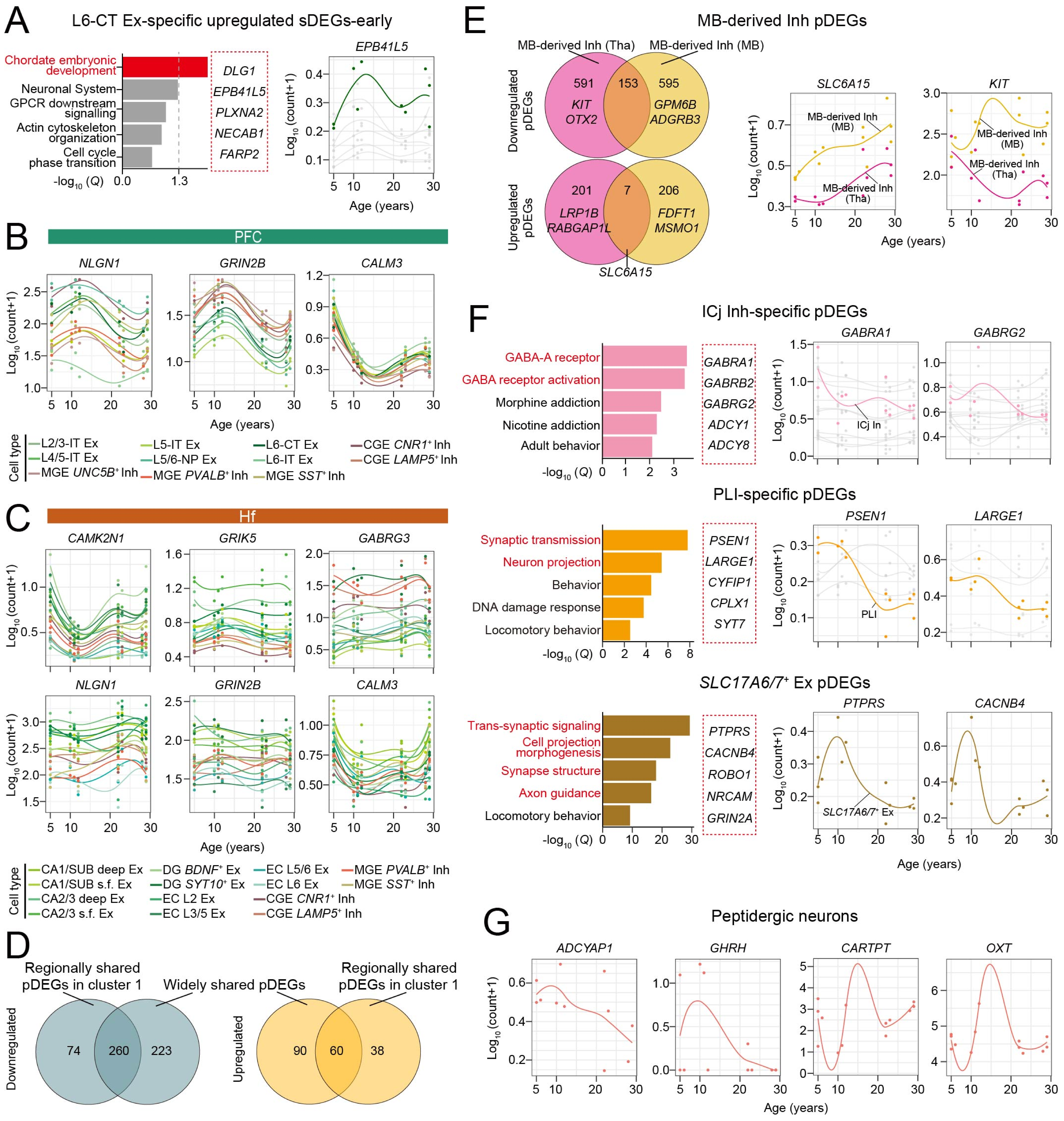

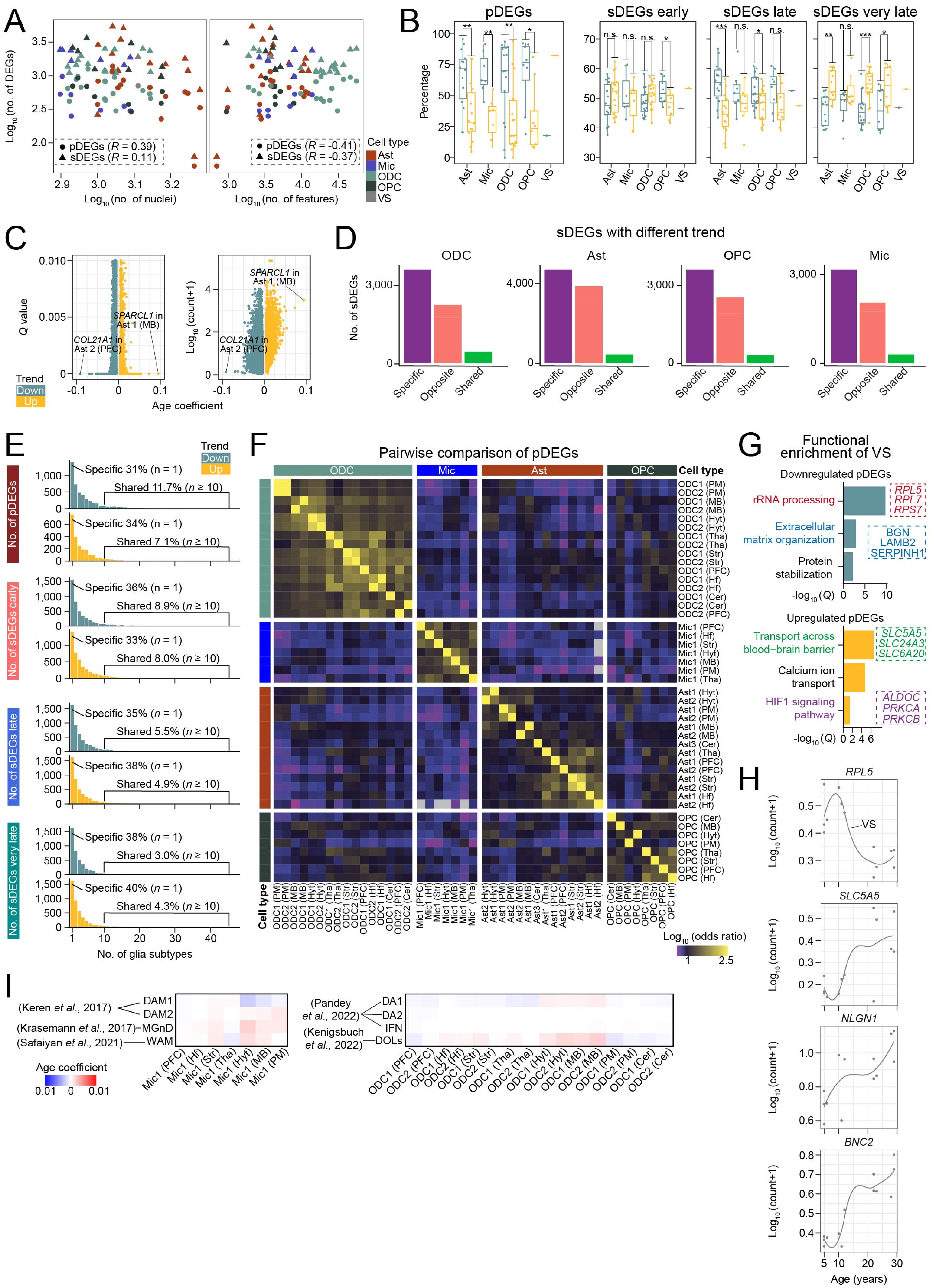

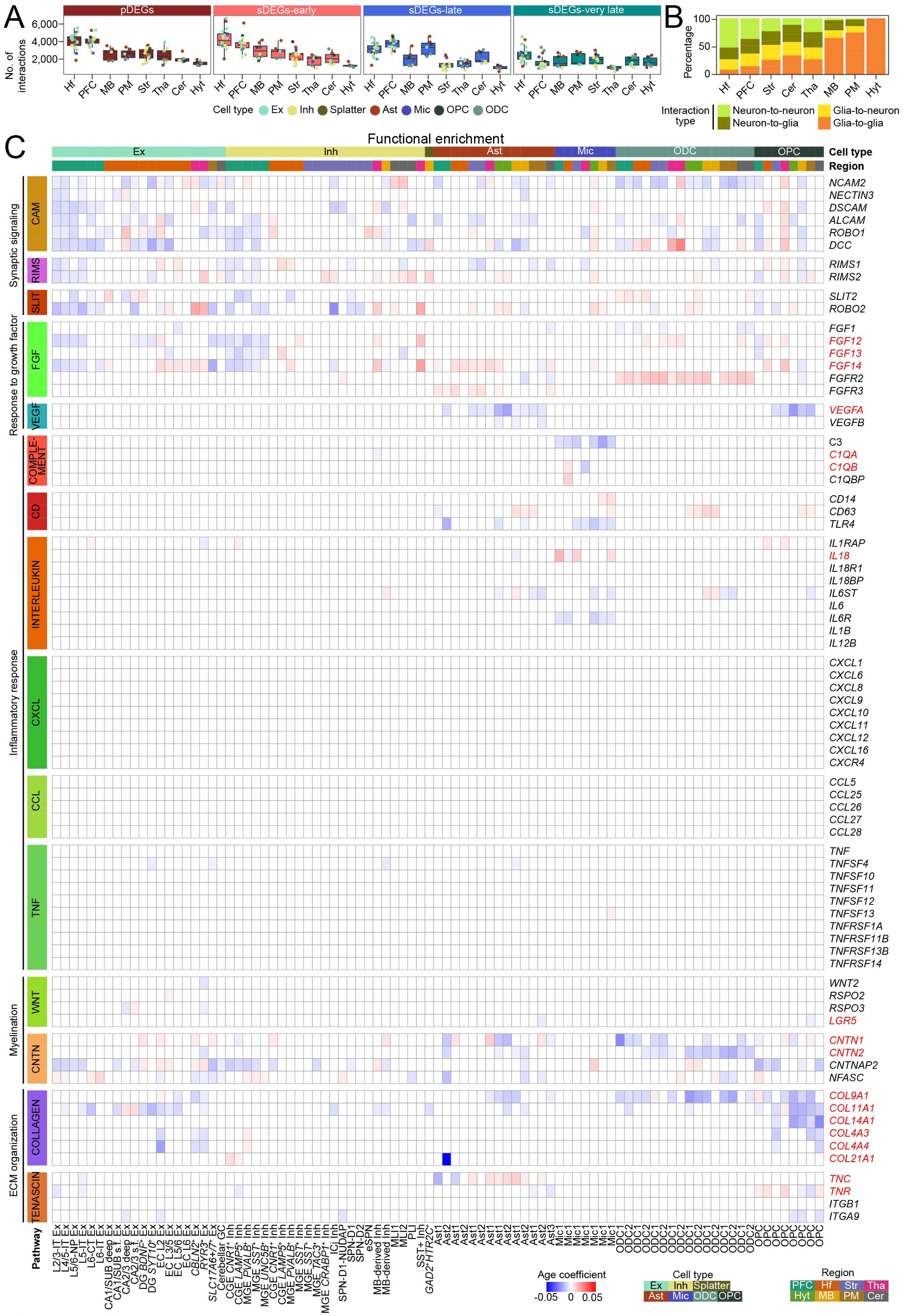

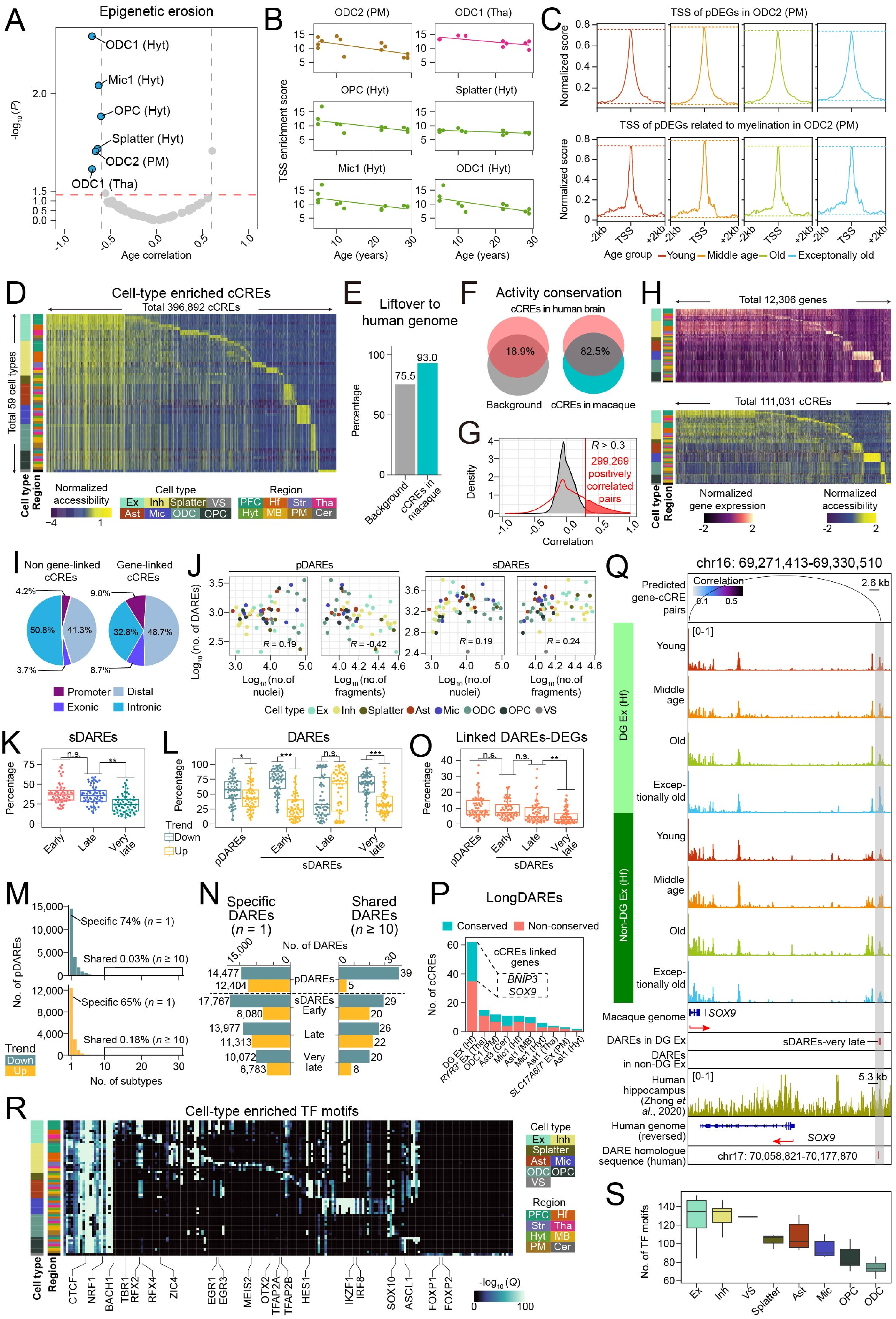

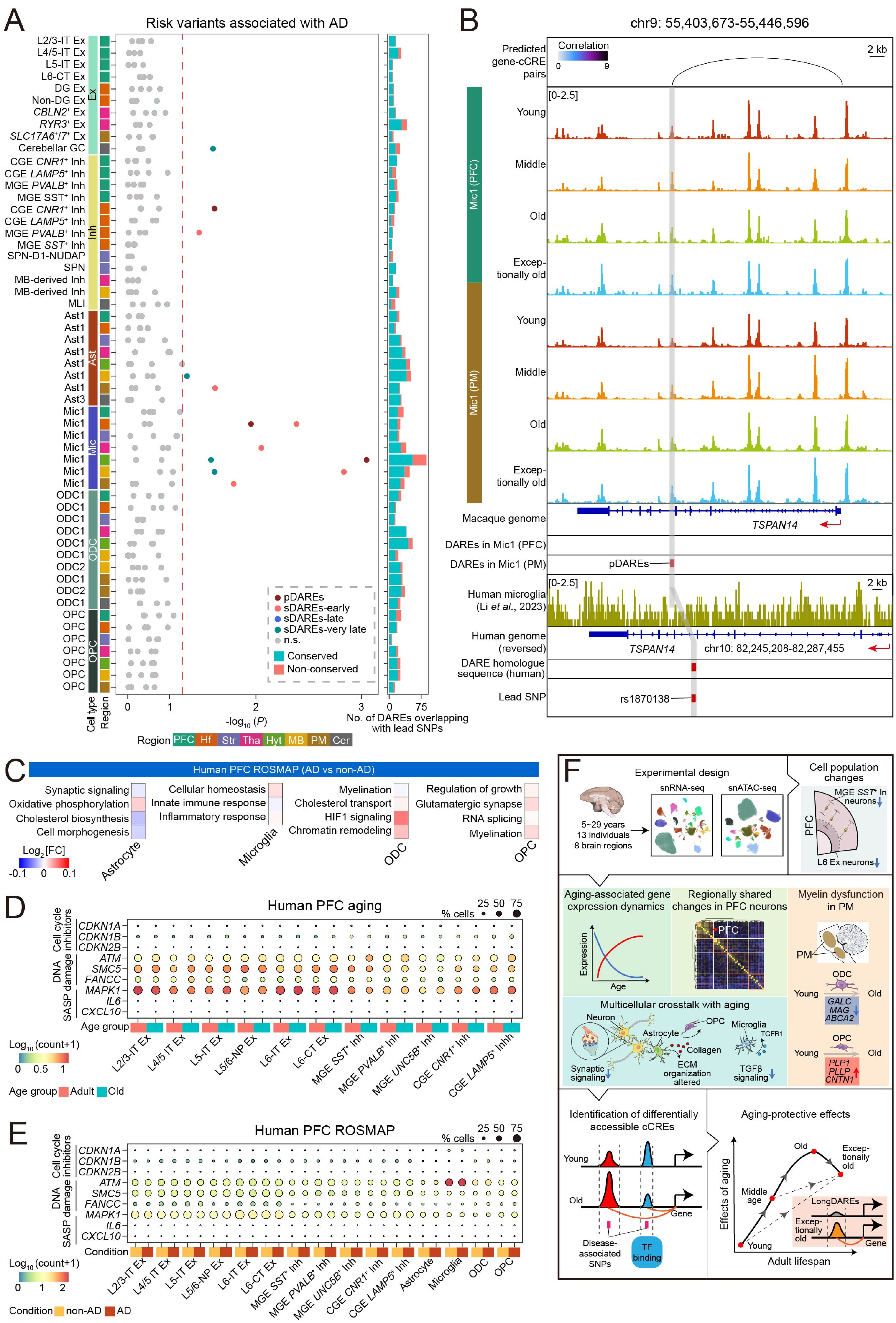
